# Nef dimerization defect abrogates HIV viremia and associated immune dysregulation in the Bone Marrow-Liver-Thymus-Spleen (BLTS) humanized mouse model

**DOI:** 10.1101/2021.01.21.427631

**Authors:** Shivkumar Biradar, Yash Agarwal, Antu Das, Sherry T. Shu, Jasmine Samal, Sara Ho, Cole Beatty, Isabella Castronova, Robbie B. Mailliard, Thomas E. Smithgall, Moses T. Bility

## Abstract

Loss of function mutations in the human immunodeficiency virus (HIV) negative factor (Nef) gene are associated with reduced viremia, robust T cell immune responses, and delayed acquired immunodeficiency syndrome (AIDS) progression in humans. *In vitro* studies have shown that mutations in the Nef dimerization interface significantly attenuate viral replication and impair host defense. However, *in vivo*, mechanistic studies on the role of Nef dimerization in HIV infection are lacking. Humanized rodents with human immune cells are robust platforms for investigating the interactions between HIV and the human immune system. The bone marrow-liver-thymus-spleen (BLTS) humanized mouse model carries human immune cells and lymphoid tissues that facilitate anti-viral immune responses. Here, we employed the BLTS-humanized mouse model to demonstrate that preventing Nef dimerization abrogates HIV viremia and the associated immune dysregulation. This suggests that Nef dimerization may be a therapeutic target for future HIV cure strategies.

## Introduction

The development of antiretroviral therapy (ART) has played a significant role in controlling the human immunodeficiency virus (HIV) epidemic. ART inhibits viral replication and prevents progression to acquired immunodeficiency syndrome (AIDS) (Deeks and Barre-Sinoussi, 2012; International et al., 2012). Unfortunately, despite ART that reduces the viral genome load to undetectable levels in the blood, HIV persists in cellular reservoirs, abrogates the anti-HIV immune response, and promotes immune dysregulation (Deeks and Barre-Sinoussi, 2012; International et al., 2012; Lisziewicz and Toke, 2013). Indeed, clinical data reveal that the HIV-accessory protein Negative factor (Nef) remains detectable in the plasma and blood cells of individuals living with ART-controlled HIV, suggesting that Nef promotes immune dysregulation during ART (Ferdin et al., 2018; Wang et al., 2015).

Individuals infected with HIV containing defects in the *nef* gene exhibit reduced viral loads and delayed AIDS progression (Learmont et al., 1999). Some individuals infected with *nef*-defective HIV exhibit a negligible viral load (Learmont et al., 1999). However, a few individuals infected with *nef*-defective HIV develop AIDS after years of naturally controlling the virus (Learmont et al., 1999). Evidence from this subset of HIV-infected individuals established Nef as a pathogenic factor in HIV infection (Dyer et al., 1999; Learmont et al., 1999). However, the mechanisms by which Nef promotes viremia and immune dysregulation *in vivo* remain unresolved (Dekaban and Dikeakos, 2017; Gorry et al., 2007).

Deleting the *nef* gene in the related simian immunodeficiency virus (SIV) reduces viral loads, preserves CD4+ T cell levels, and induces an effective anti-SIV immune response in monkeys (Daniel et al., 1992; Gabriel et al., 2016). Indeed, the gold standard for vaccine-induced immunity against HIV/AIDS is the *nef*-deleted SIV-wild-type SIV challenge model in macaques (Gabriel et al., 2016). Although the SIV-macaque model has provided insights on the role of Nef in modulating HIV-immune system interactions (Hu, 2005), important differences exist between SIV and HIV (Feinberg and Moore, 2002; Lifson and Haigwood, 2012; Pollom et al., 2013) and between human and monkey immune cell signaling (Shedlock et al., 2009). Therefore, rodent models hosting human immune cells and lymphoid tissues termed humanized rodent models provide a means to investigate HIV-human immune cell interactions (Agarwal et al., 2020; Akkina et al., 2020; Samal et al., 2018).

Humanized mouse models are reconstituted with varying types of human immune cells and lymphoid tissues, resulting in varying degrees of immune function. Humanized mice reconstituted using peripheral blood lymphocytes (PBL; PBL-humanized mice) are first-generation models that support HIV replication, and those mice are repopulated with activated-CD4+ T cells and marginal levels of other immune cells (Skelton et al., 2018). The second-generation humanized mouse models are reconstituted using human hematopoietic stem cells (HSC-humanized mice), enabling the development of *de novo* immune cells, which can mediate antigen-specific immune responses to HIV infection (Skelton et al., 2018). The third-generation humanized mouse models are reconstituted using human hematopoietic stem cells and lymphoid tissues, which are required to generate a robust antigen-specific immune response to HIV infection (Skelton et al., 2018). *In vivo* studies in the bone marrow-liver-thymus (BLT) humanized mouse model, a third-generation humanized mouse model, demonstrated that *nef* deletion delays viremia kinetics for 1-2 weeks and reduces CD4+ T cell depletion at low-dose inoculum compared to wild-type HIV (Zou et al., 2012). However, *nef*-deleted HIV and wild-type HIV exhibit similar replication kinetics in BLT-humanized mice following high-dose inocula (Watkins et al., 2013; Zou et al., 2012).

Tissue culture models using human immune cells have complemented *in vivo* models and provided invaluable insights on Nef-human immune cell interactions in HIV infection. Tissue culture studies demonstrate that Nef downregulates major histocompatibility complex class I (MHC-I) molecules in HIV infected CD4+ T cells (Chang et al., 2001; Dirk et al., 2016; Hung et al., 2007; Roeth et al., 2004) to impair CD8+ cytotoxic T cell-mediated killing of HIV infected cells (Gorry et al., 2007). Additionally, tissue culture studies demonstrate that Nef promotes B cell dysregulation via impairment of CD4+ T cell-B cell interactions (Kaw et al., 2020). We demonstrated that Nef homodimerization is crucial for Nef-mediated enhancement of HIV infectivity and replication as well as dysregulation of host defenses in tissue culture models (Li et al., 2020; Poe and Smithgall, 2009; Rosa et al., 2015; Ryan P. Staudt, 2020; Staudt and Smithgall, 2020; Trible et al., 2006a). We also demonstrated that Nef dimers activate non-receptor tyrosine kinases, which serve as effectors for Nef signaling to enhance the viral life cycle (Li et al., 2020; Tarafdar et al., 2014; Trible et al., 2006b). Additionally, small molecules believed to affect Nef dimerization promote redisplay of cell-surface MHC-I and promote CD8 T cell-mediated killing of latently HIV-infected cells in tissue culture (Emert-Sedlak et al., 2013; Mujib et al., 2017).

In this study, we employed the bone marrow-liver-thymus-spleen (BLTS) humanized mouse model, which incorporates human innate and adaptive immune cells as well as primary and secondary lymphoid tissues, to investigate HIV Nef dimer-human immune system interactions. First, we show that deletion of Nef allows complete control of HIV infection in BLTS mice. Then, using a series of dimerization-defective Nef mutants based on previous X-ray crystal structures (reviewed by Staudt, *et al*.) (Staudt et al., 2020), we demonstrate that Nef dimerization is essential for high-titer HIV replication *in vivo*. Furthermore, we show that Nef dimers are required for enhanced T cell-mediated immune activation, checkpoint inhibitor expression and dysregulation of many other immune signaling pathways *in vivo*.

## Results

### The BLTS-humanized mouse model supports functional human innate and adaptive immune cells in the blood and the human lymphoid tissues

We previously reported human immune cell development in the peripheral blood and the human primary and secondary lymphoid tissue grafts in the BLTS-humanized mouse model (Watkins et al., 2013). Here, we demonstrate the reconstitution of additional human immune cell types in peripheral blood and human primary (thymus) and secondary (spleen) lymphoid tissue grafts. Specifically, we demonstrate reconstitution with NK cells, CD4+ and CD8+ T cells, monocytes, memory B cells, plasmablasts, and pan-T cells, pan-B cells, and macrophages as previously demonstrated (Supplementary Figure 1-3) (Samal et al., 2018).

Additionally, we demonstrate the functionality of human-antigen presenting cells in the BLTS-humanized mouse model (Supplementary Figure 4A, 4B). Bone marrow-derived hematopoietic stem cells in the BLTS-humanized mice can differentiate into type-1 and type-2-polarized dendritic cells (DCs) (Supplementary Figure 4A). Type-1 DCs secrete the T-helper 1 (Th1)-driving cytokine (IL-12p70) upon stimulation with physiological ligand (CD40L) (Supplementary Figure 4A). Human CD163+ splenic macrophages from BLTS-humanized mice phagocytized bacteria (pHrodo™ Red *E. coli* BioParticles) upon co-culture (Supplementary Figure 4B). Furthermore, we demonstrate the functionality of T cells in the BLTS-humanized mouse model. We show robust splenic T cell response (type-1 cytokine, IFNγ) to physiological stimulation (CD3 and CD28) that was comparable to the cytokine (IFNγ) response by T cells from human PBMCs (Supplementary Figure 4C, 4D).

### The human immune system in the BLTS mouse model abrogates *nef*-deleted HIV viremia

We previously reported that BLTS-humanized mice support HIV replication and associated immunopathogenesis (Samal et al., 2018). We demonstrate that human macrophages (CD163+ splenic cells) and CD4+ T cells facilitate HIV replication (Supplementary Figure 5) (Samal et al., 2018). Therefore, BLTS-humanized mice provide a robust *in vivo* model for investigating Nef-immune system interactions in HIV infection. Nef enhances HIV infectivity and replication in some cell lines (Crotti et al., 2006; Spina et al., 1994; Yang et al., 2002). Nef also enhances HIV replication in human PBL-derived activated-primary CD4+ T cells in a viral input-dependent manner, with the Nef effect lost at a high viral-input dose (Shi et al., 2020). Here we show that *nef*-deleted HIV (NL4-3 strain) exhibits reduced replication compared to wild-type HIV in the TZM-bl cell line (Figure 1A). However, *nef*-deleted HIV and wild-type HIV exhibit negligible differences in viral replication at high viral input in the activated-primary CD4+ T cell culture model. (Figure 1B). The *in vivo* equivalent of the activated-primary CD4+ T cell culture model is the PBL-humanized mouse model (Agarwal et al., 2020; Denton and Garcia, 2011). Using the PBL-humanized mouse model, we demonstrate that *nef*-deleted HIV and wild-type HIV exhibit similar infectivity and replication in the blood and lymphoid tissues (Figure 1C, 1D, 1E), which is consistent with previous studies (Gulizia et al., 1997). On the contrary, inoculation of BLTS-humanized mice with *nef*-deleted HIV and wild-type HIV resulted in divergent outcomes, with viremia in wild-type HIV inoculated mice and no viremia in *nef*-deleted HIV-inoculated mice (Figure 1C, 1F). We inoculated BLTS-humanized mice with the same high dose of *nef*-deleted HIV and wild-type HIV stocks used to infect PBL-humanized mice; therefore, the *nef* deleted HIV inoculum is infectious and replication-competent (Figure 1C). These observations demonstrate that Nef overrides host immune control of HIV-1 replication in the BLTS-humanized mouse model.

**Figure 1.**
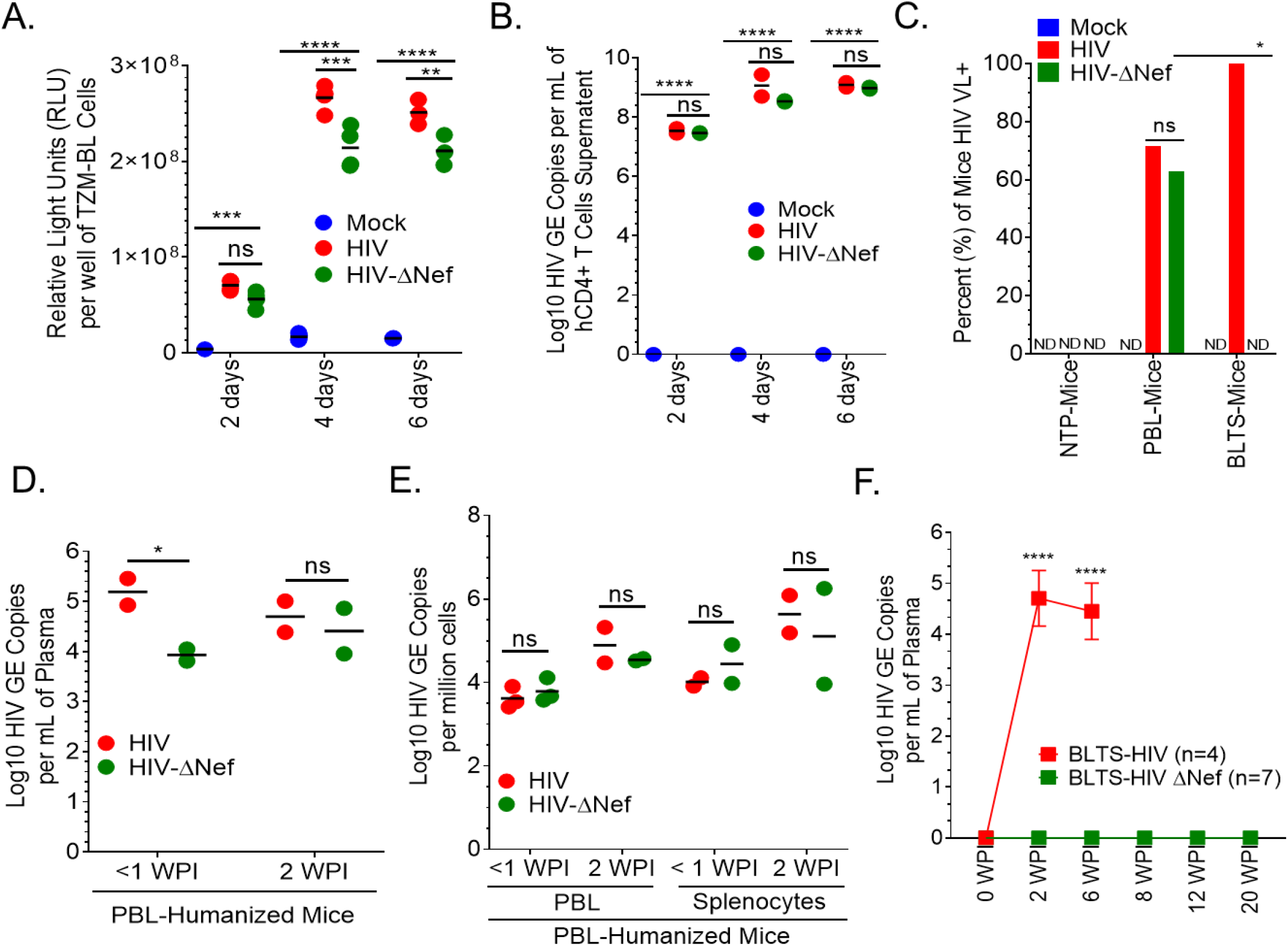
The human immune system in the BLTS mouse model controls *nef*-deleted HIV viremia. Analysis of the infectivity and replication of wild-type HIV and *nef*-deleted HIV (0.1 ng HIV p24 per 1×10^4 cells) demonstrate (A) reduced infectivity and replication kinetics in TZM-BL cells (N=3-4 wells per group); on the contrary, (B) similar infectivity and replication kinetics were recorded in activated (CD3/CD28)-primary CD4+ T cells (N=2 wells per group). (C-E) The infectivity and replication of wild-type and *nef*-deleted HIV (10 ng HIV p24 inoculum per mouse) in peripheral blood lymphocytes (PBL; reconstituted with predominately human CD4+ T cells) humanized mice (PBL-humanized mice; N=7-8 mice per group, and 3-5 per timepoint) is comparable as measured in the blood (plasma and PBL) and splenocytes at indicated weeks post-infection (WPI). (C) For additional controls, non-transplanted NSG mice (NTP) were inoculated with the different HIV strains and mock to demonstrate that HIV replication (i.e., viral load) requires the presence of human cells. (C) For negative controls, the absence of viral load was demonstrated in the blood and splenocytes of mock-inoculated PBL-humanized mice (*data not shown*). (F) On the contrary, BLTS-humanized mice (N=4-7 mice per group; 10 ng HIVp24 inoculum per mouse) exhibit complete inhibition of *nef*-deleted HIV viremia. For BLTS-humanized mice inoculated with *nef*-deleted HIV, 7 mice were evaluated up to 6 weeks post-infection (WPI), and only 2 mice and 1 mouse remained in the study at 8-12 WPI and at 20 WPI, respectively. Mock inoculated BLTS-humanized mice (n=4) were used for controls, with those mice showing no detectable HIV genome (C; *timepoints data not shown*). Not significant (ns) = P>0.05, * = P≤ 0.05, ** = P≤ 0.01, *** = P≤ 0.001, **** = P≤ 0.0001.

HIV transmission in humans results in a cytokine burst (including IFNγ, IL-10, and TNFα) within two weeks of exposure (McMichael et al., 2010), which stimulates immune cells to initiate an anti-viral immune response (Stacey et al., 2009). Inoculation of BLTS-humanized mice with *nef*-deleted HIV induces a modest increase in the secretion of type 1 cytokines (IFNγ, IL-2, TNFα, IL-12p70) and type 2 cytokines (IL-10, IL-6, IL-8, IL-13) as compared to the highly viremic wild-type HIV (Supplementary Figure 6). The cytokine burst in wild-type HIV infection does not eliminate the virus or reduce viremia to undetectable levels; instead, it may fuel immune dysregulation (McMichael et al., 2010). The resulting chronic HIV infection induces progressive T cell-immune activation and checkpoint inhibitor expression (Kahan et al., 2015; Khaitan and Unutmaz, 2011; McMichael et al., 2010), inadequate anti-viral T cell responses (Kahan et al., 2015; Khaitan and Unutmaz, 2011; McMichael et al., 2010), and immune dysregulation (Boasso et al., 2009; Chirmule et al., 1994).

### Nef dimerization defect abrogates HIV viremia and associated immunodeficiency in the BLTS-humanized mouse model

Previous X-ray crystallography studies of HIV-1 Nef proteins either alone or in complexes with host cell kinase regulatory domains revealed that Nef forms homodimers (reviewed in Staudt et al., 2020). Comparison of these structures identified several residues common to these Nef dimer interfaces, including L112, Y115, and F121. Mutagenesis of these residues prevents homodimerization of recombinant Nef *in vitro* and reduces Nef dimer formation in a cell-based bimolecular fluorescence complementation (BiFC) assay. (Poe and Smithgall, 2009; Poe et al., 2014). Based on the BiFC assay, the relative disruption of dimerization in the Nef mutants is: L112D+Y115D double mutant > Y115D > F121A (Li et al., 2020; Poe and Smithgall, 2009). Previous studies showed that these dimerization-defective Nef mutants also reduce HIV replication (Li et al., 2020; Poe and Smithgall, 2009) and abrogate HIV-induced impairment of host defense (Ryan P. Staudt, 2020) in tissue culture models, implicating Nef dimers in signaling mechanisms by which HIV-Nef promotes viremia and immune dysregulation (Poe and Smithgall, 2009). Interestingly, we show that the *nef*-dimerization defective HIV strains, Y115D, and L112D+Y115D are replication-competent in PBL-humanized mice and the HSC-humanized mice (Supplementary Figure 7), respectively. However, BLTS-humanized mice inoculated with the *nef* dimerization-defective HIV strains Y115D and Y115D + L112D were aviremic (Figure 2, Supplementary Figure 8) and demonstrated reduced HIV-infected T cells in human spleen xenografts when compared to wild-type HIV infection (Figure 3A, 3B, Supplementary Figure 9). Additionally, mice inoculated with HIV carrying the Nef-F121A mutant were partly viremic (67% of mice) (Supplementary Figure 8). BLTS-humanized mice inoculated with *nef*-deleted HIV (NL4-3 strain) were predominately aviremic (75% of the mice) and had low levels of HIV-infected T cells in the human spleen xenografts (Figure 2, 3A, 3B). Additionally, *nef*-defective HIV (ΔNef and the dimerization-defective mutant, Y115D) and mock-inoculated BLTS-humanized mice exhibited similar CD4+/CD8+ T cell ratios, whereas wild-type HIV-infected BLTS-humanized mice exhibited immunodeficiency (reduced CD4+/CD8+ T cell ratio; CD4+ T cell loss) (Figure 4A, 4B).

**Figure 2.**
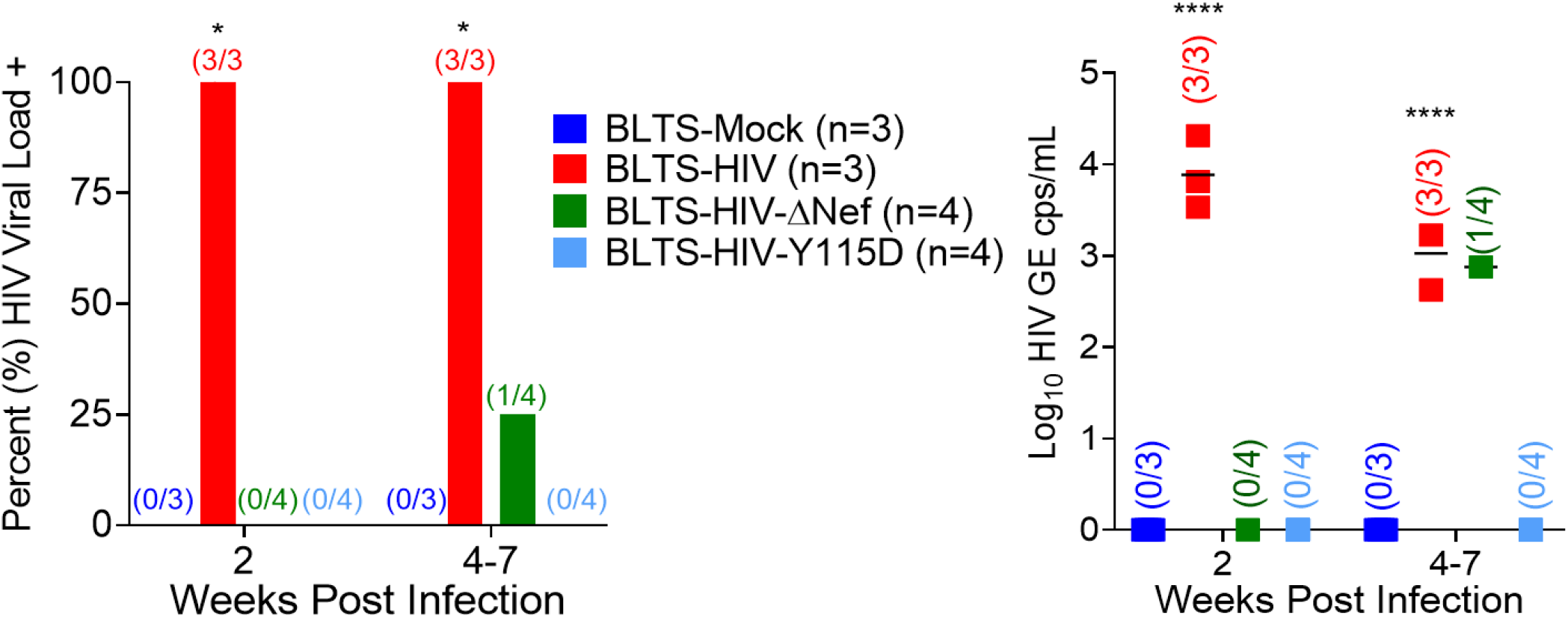
Nef-dimerization defect abrogates HIV viremia in BLTS-humanized mice. BLTS-humanized mice (BLTS-Mice; N=3-4) exhibit viremic control of *nef*-dimerization defective-HIV. N=3-4 mice per group, with mock (2 mice at 7 weeks post-inoculation and 1 mouse at 4 weeks post-inoculation), wild-type HIV (3 mice at 7 weeks post-infection), Nef-deletion-ΔNef (2 mice at 7 weeks post-inoculation and 2 mice at 4 weeks post-inoculation) and Nef dimerization-defective-Y115D (2 mice at 7 weeks post-inoculation and 1 mouse at 6 weeks post-inoculation) for indicated 4-7 weeks timepoints. Not significant (ns) = P>0.05, * = P≤ 0.05, ** = P≤ 0.01, *** = P≤ 0.001, **** = P≤ 0.0001.

**Figure 3.**
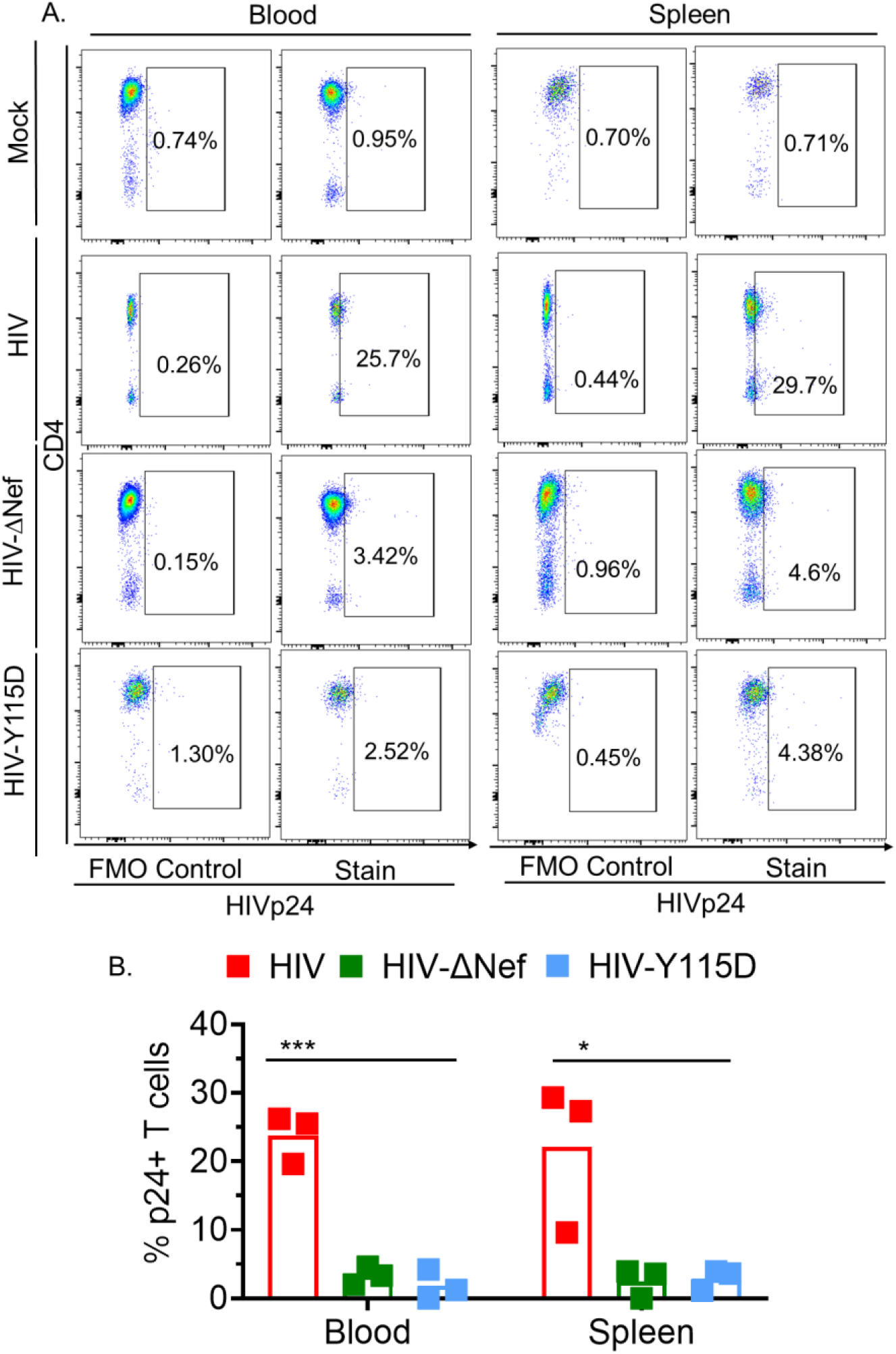
Nef-dimerization defect reduces HIV infected-T cells in BLTS-humanized mice. (A-B) Inoculation of BLTS-humanized mice with a *nef*-dimerization defective HIV (Y115D; 10 ng HIVp24) results in reduced HIVp24+ CD3+ cells (HIV infected-T cells) in the human spleen, as compared to wild-type HIV infection, as measured using flow cytometry. Mock inoculated mouse was used as a control in flow cytometry analysis. N=3 mice per group, with wild-type (3 mice at 7 weeks post-infection), Nef-deletion-ΔNef (2 mice at 7 weeks post-inoculation and 1 mouse at 4 weeks post-inoculation) and Nef dimerization-defective-Y115D (2 mice at 7 weeks post-inoculation and 1 mouse at 6 weeks post-inoculation) for the 4-7 weeks sacrifice-timepoints. Not significant (ns) = P>0.05, * = P≤ 0.05, ** = P≤ 0.01, *** = P≤ 0.001, **** = P≤ 0.0001.

**Figure 4.**
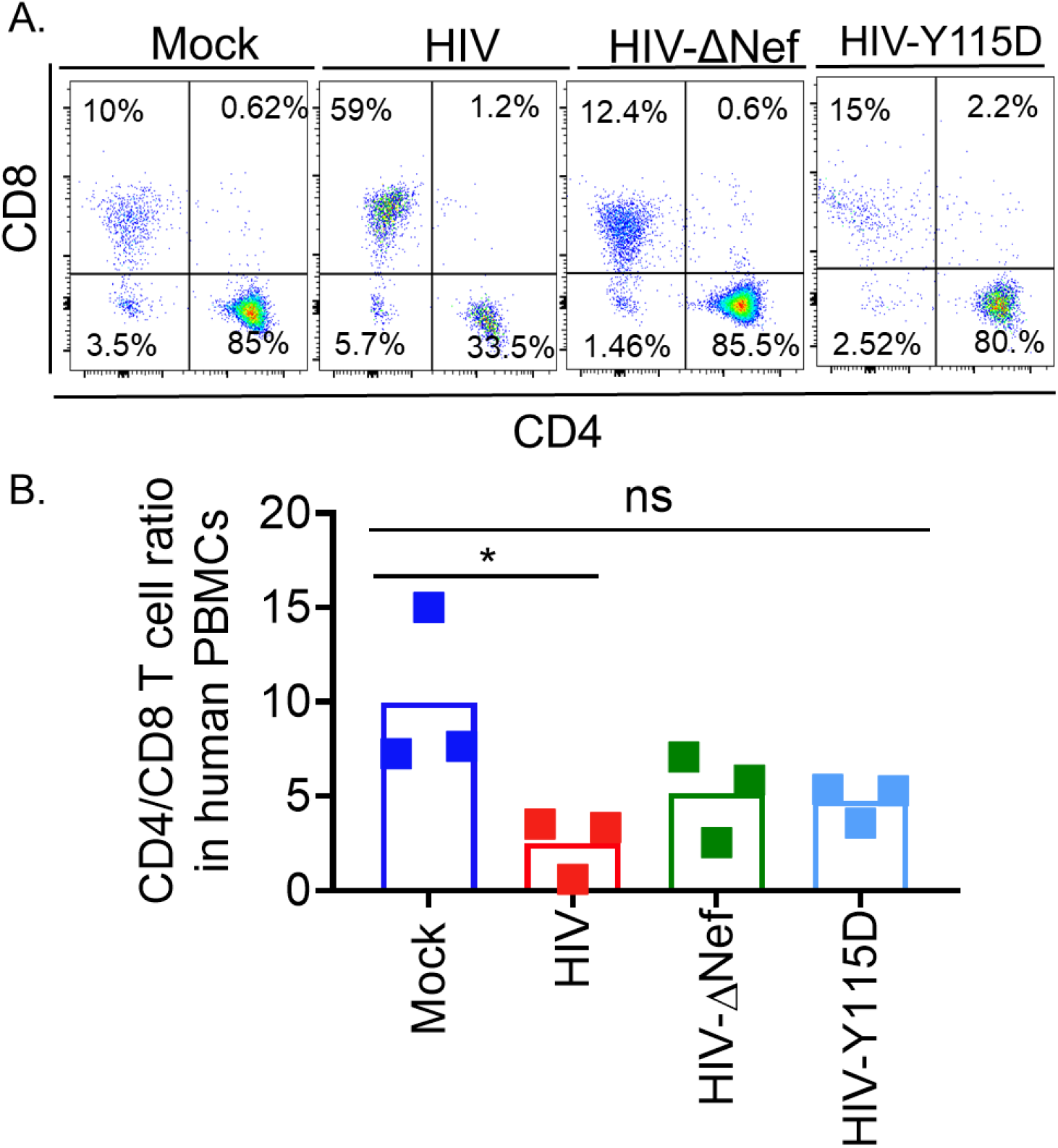
HIV-*nef* dimerization defect abrogates HIV-induced CD4+ T cell depletion in the blood in BLTS-humanized mice. Flow cytometry analysis of human CD4+ and CD8+ T cells (CD4+/CD8+ T cell ratio) in the blood in mock, wild-type HIV, and *nef*-defective HIV infection (*nef*-deleted-ΔNef, and *n*ef-dimerization defective-Y115D; 10 ng HIVp24) in BLTS-humanized mice. N=3 mice per group, with mock (2 mice at 7 weeks post-inoculation and 1 mouse at 4 weeks post-inoculation), wild-type HIV (3 mice at 7 weeks post-infection), *nef*-deleted-ΔNef (2 mice at 7 weeks post-inoculation and 2 mice at 4 weeks post-inoculation) and *nef*-dimerization defective-Y115D (2 mice at 7 weeks post-inoculation and 1 mouse at 6 weeks post-inoculation) for indicated timepoints. ns = P>0.05, * = P≤ 0.05.

### Nef dimerization defect attenuates HIV-induced immune dysregulation in the BLTS-humanized mouse model

Many studies have shown that Nef impairs the anti-viral immune response to HIV infection (Pawlak and Dikeakos, 2015). Furthermore, several lines of evidence demonstrate that Nef-dimerization defective mutants and Nef inhibitors abrogate HIV-induced immune dysregulation (Dekaban and Dikeakos, 2017; Mujib et al., 2017). We observed that Nef dimerization defect (Y115D mutant) attenuates HIV-induced T cell-immune activation (elevated CD25+ and HLA-DR+ levels) (Figure 5, Supplementary Figure 10A) and checkpoint inhibitor expression (elevated PD1+ levels) (Figure 6, Supplementary Figure 10B) in the blood and human spleen. Consistent with the role of Nef in promoting HIV-induced immune dysregulation, complete deletion of the *nef* gene in the HIV genome attenuates HIV-induced T cell-immune activation markers (elevated CD25+ and HLA-DR+ levels) and checkpoint inhibitor expression (high PD1+ levels) in the blood and the human spleen in BLTS-humanized mice (Figure 5-6, Supplementary Figure 10).

**Figure 5.**
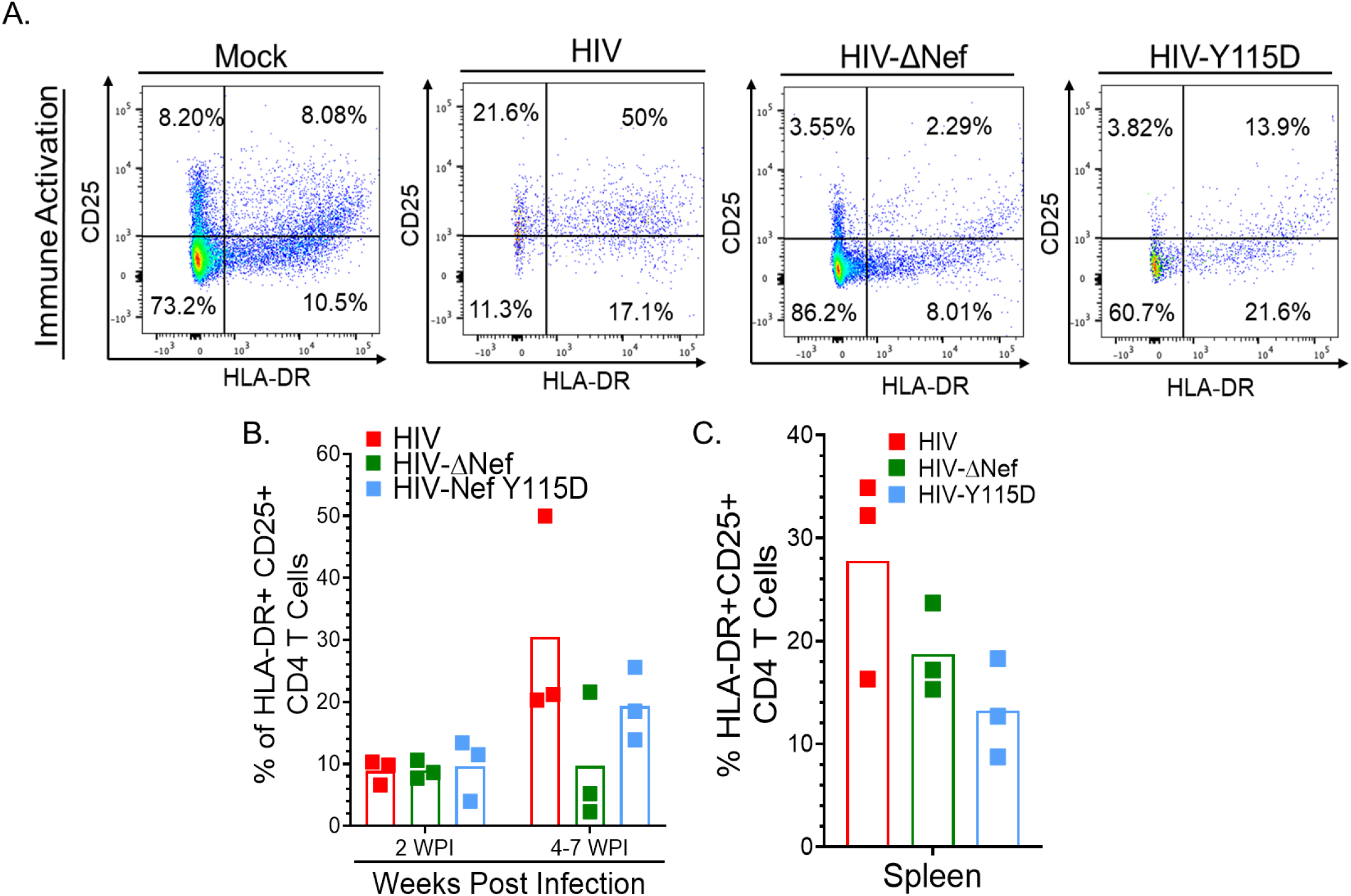
Nef dimerization defect abrogates HIV-induced elevated HLADR+ and CD25+ CD4+ T cells in the blood and human spleen in BLTS-humanized mice. (A, B) Flow cytometry analysis of human CD4+ T cells in the blood in BLTS-humanized mice demonstrates HIV-induced HLA-DR+ and CD25+ CD4+ T cells are reduced in *nef*-defective HIV infection (*nef*-deleted-ΔNef, and *nef*-dimerization defective-Y115D; 10 ng HIVp24 per mouse). Human CD4+ T cells from Mock-BLTS-humanized mouse served as a control. (C) Flow cytometry analysis of human CD4+ T cells in the human spleen xenografts in BLTS-humanized mice demonstrates reduced HLADR+ and CD25+ CD4+ T cells in *nef*-defective HIV infection (*nef*-deleted-ΔNef, and *nef*-dimerization defective-Y115D) compared to wild-type HIV. N=3 per group, with wild-type HIV (3 mice at 7 weeks post-infection), *nef*-deleted-ΔNef (2 mice at 7 weeks post-inoculation and 1 mouse at 4 weeks post-inoculation) and *nef*-dimerization defective-Y115D (2 mice at 7 weeks post-inoculation and 1 mouse at 6 weeks post-inoculation) at indicated 4-7 weeks sacrifice-timepoints.

**Figure 6.**
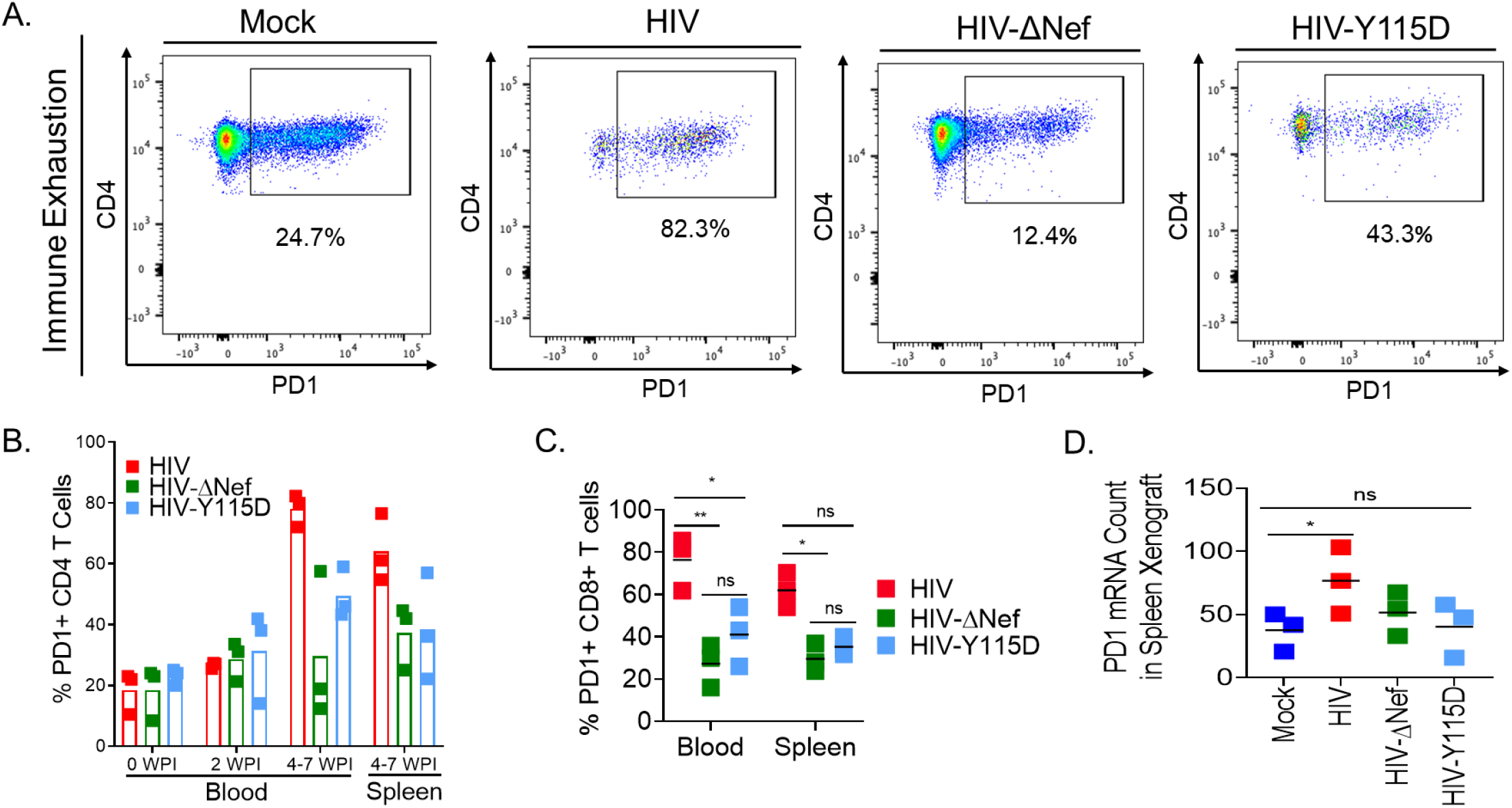
Nef dimerization defect abrogates HIV-induced T cell-checkpoint inhibitor expression in the blood and human spleen in BLTS-humanized mice. (A, B, C) Flow cytometry analysis of human CD4+ and CD8+ T cells in the blood and human spleen in BLTS-humanized mice demonstrates elevated CD4+ and CD8+ T cell-checkpoint inhibitor expression (PD1+) in the wild-type HIV group compared to *nef*-defective HIV groups (*nef*-deleted-ΔNef, and *nef*-dimerization defective-Y115D; 10 ng HIVp24 per mouse). (A) Human CD4+ T cells from Mock-BLTS-humanized mouse served as a control. (D) Additionally, gene expression analysis of the human secondary lymphoid tissue xenograft (Spleen) in BLTS-humanized mice demonstrates elevated expression of PD1 in the wild-type HIV group compared *nef*-defective HIV groups (*nef*-deleted-ΔNef, and *nef*-dimerization defective-Y115D) and mock. Not significant (ns) = P>0.05, * = P≤ 0.05, ** = P≤ 0.01. N=3 per group, with wild-type HIV (3 mice at 7 weeks post-infection), *nef*-deleted-ΔNef (2 mice at 7 weeks post-inoculation and 1 mouse at 4 weeks post-inoculation) and *nef*-dimerization defective-Y115D (2 mice at 7 weeks post-inoculation and 1 mouse at 6 weeks post-inoculation) at indicated 4-7 weeks sacrifice-timepoints.

The current consensus posits that chronic HIV infection in humans results from an inadequate antiviral-Th1 immune response, which is associated with T cell-checkpoint inhibitor expression (Khaitan and Unutmaz, 2011), systemic immune inflammation (i.e., elevated levels of CXCL13) (Mehraj et al., 2019), B cell hyper-activation and dysregulation (Chirmule et al., 1994; Kaw et al., 2020), and reduced levels of anti-viral factors (i.e., CXCL12) (Struyf et al., 2009). Previous studies also show that HIV Nef dimers are required to activate non-receptor tyrosine kinases of the Src and Tec families to enhance viral replication (Staudt et al. JBC 2020) and are also linked to downregulation of MHC-I and CD4 receptor on T cells (Poe and Smithgall, 2009; Shu et al., 2017). Additionally, Nef promotes B cell hyper-activation and dysregulation; this pathogenic effect has been associated with T cell interaction (Chirmule et al., 1994; Kaw et al., 2020). Dysregulated immune activation and signaling by Nef are presumed to drive chronic HIV infection and progression to AIDS (Lori, 2008). Gene expression analysis demonstrates that wild-type HIV infection in BLTS-humanized mice induces systemic inflammation marker (elevated CXCL13), B cell activation markers (elevated CD19, CD79, PAX5, and BLNK), and reduced levels of anti-viral factors (CXCL12, PML/TRIM19) in the human spleen xenograft (Figure 7A). Wild-type HIV infection also induced the expression of the non-receptor tyrosine kinase, ABL1, in the human spleen xenografts of BLTS-humanized mice (Figure 7A). ABL1 plays a role in T cell receptor signaling and cytoskeletal rearrangement, facilitating HIV entry into the host cell (Gu et al., 2009). ABL1 also enhances HIV-1 replication by activating RNA pol-II (McCarthy et al., 2019). Comparative analysis of gene expression in BLTS-humanized mice inoculated with wild-type HIV and *nef*-deleted HIV showed reduced expression of the non-receptor tyrosine kinase, SYK, reduced systemic inflammation (CXCL13, HLA-DR), decreased B cell activation markers (CD19, AICDA), and elevated levels of anti-viral factors (CXCL12, LIF) in the human spleen xenografts in *nef*-deleted HIV infection compared to wild-type HIV infection (Figure 7B). A similar analysis of the human spleen xenograft revealed reduced systemic inflammation (CXCL13); elevated levels of anti-viral factors (CXCL12, IFIT2); upregulation of genes involved in complement activation (C1Q); increased expression of RAG1/2 for the development of a diverse repertoire of immunoglobulins; and elevated levels of the MHC-related molecule CD1A in BLTS-humanized mice challenged with the *nef*-dimerization mutant (Y115D) as compared to mice challenged with wild-type HIV (Figure 7C).

**Figure 7.**
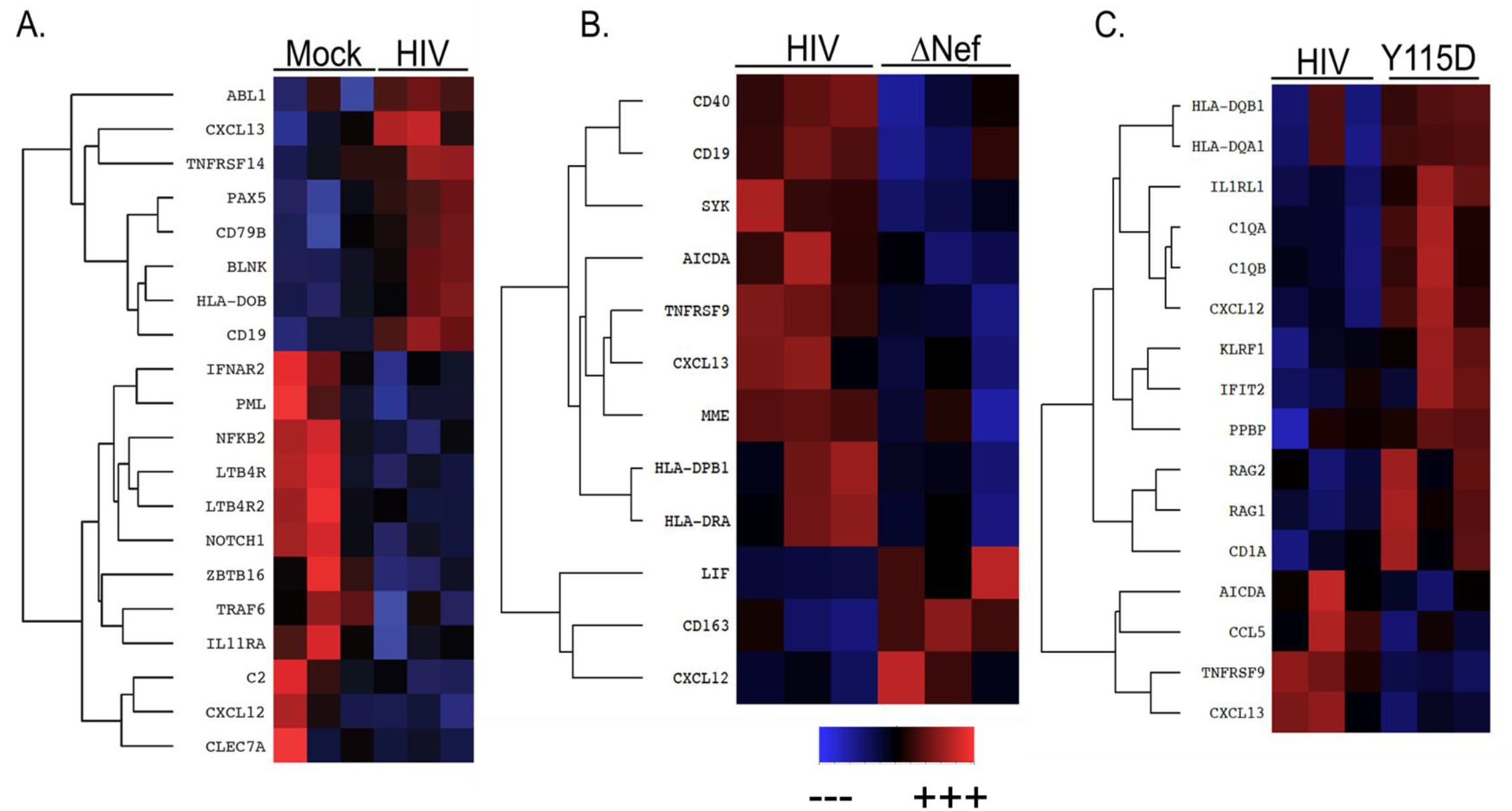
Nef dimerization defect abrogates HIV-induced immune dysregulation in the human spleen in BLTS-humanized mice. (A-C) Nanostring analysis of the human spleen in BLTS-humanized mice demonstrates elevated expression of systemic immune activation, B cell hyper-activation and non-receptor tyrosine kinase genes and reduced anti-viral inhibitors genes in Wildtype HIV infection compared to (A) mock and (B-C) *nef*-defective HIV infection (*nef*-deleted-ΔNef, and *nef*-dimerization defective-Y115D; 10 ng HIVp24 per mouse) and mock inoculation. Note: blue denotes downregulation and red denote upregulation. N=3 per group, with wild-type HIV (3 mice at 7 weeks post-infection), *nef*-deleted-ΔNef (2 mice at 7 weeks post-inoculation and 1 mouse at 4 weeks post-inoculation) and *nef*-dimerization defective-Y115D (2 mice at 7 weeks post-inoculation and 1 mouse at 6 weeks post-inoculation) at 4-7 weeks sacrifice-timepoints.

Consistent with the gene expression profile in the human spleen in BLTS-humanized mice, Ingenuity pathway analysis (IPA) demonstrates elevated systemic immune activation signaling (Th1 pathway, Neuroinflammatory pathway, Induction of apoptosis by HIV) and B cell signaling (PI3K signaling, FcγRIIB signaling, B cell receptor signaling) in HIV infection (Figure 8A). It also revealed reduced anti-viral immune signaling, specifically Toll-like receptor (TLR) signaling, HMGB1 signaling, NK cell signaling, and RIG1-like receptors, all of which are downregulated in wild-type HIV infection compared to mock (Figure 8A). Additionally, IPA demonstrates elevated anti-viral immune signaling (Toll-like receptor signaling, HMGB1 signaling, complement system signaling, NK cell signaling, DC maturation, and NK-DC crosstalk, RIG1-like receptors and pattern recognition receptors (PRR) signaling) and enhanced Th1 signaling and decreased B cell signaling (PI3K signaling, FcγRIIB signaling, B cell receptor signaling) in *nef*-defective HIV infection cohorts (ΔNef and dimerization-defective Nef-Y115D) compared to wild-type HIV cohorts (Figure 8B, 8C). This suggests that components of the innate immune system in the human spleen xenografts such as complement system, PRRs, HMGB1, NK cells, dendritic cells, and macrophages are playing an essential role in controlling viremia in *nef*-defective HIV infected humanized mice. Furthermore, deletion or mutation of *nef* abrogates the ability of HIV to interact with host proteins to downregulate MHC class-I, which allows CTLs to recognize infected cells in the context of MHC class-I and induce cytotoxicity to control viremia in BLTS-humanized mice.

**Figure 8.**
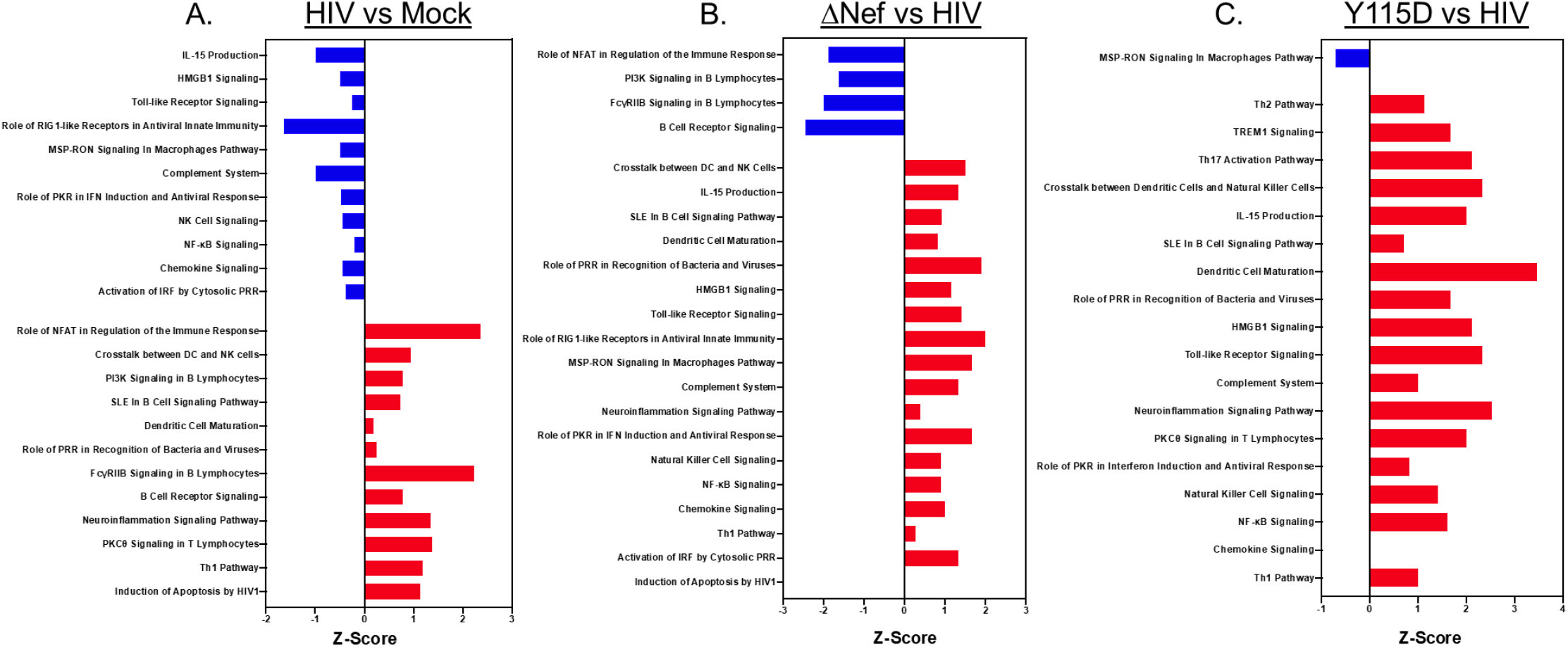
Nef dimerization defect abrogates HIV-induced immune dysregulatory signaling in human spleen in BLTS-humanized mice. (A-C) Ingenuity Pathway Analysis of the Nanostring gene expression data demonstrates elevated systemic immune activation signaling and B cell signaling and reduced anti-viral immune signaling in wildtype HIV infection compared to mock (A). Ingenuity Pathway Analysis also demonstrates elevated anti-viral immune signaling and Th1 signaling in *nef*-defective HIV infection (*nef*-deleted-ΔNef, and *nef*-dimerization defective-Y115D; 10 ng HIVp24 per mouse) compared to wild-type HIV (B, C). Note: blue denotes downregulation and red denote upregulation. N=3 per group, with wild-type HIV (3 mice at 7 weeks post-infection), *nef*-deleted-ΔNef (2 mice at 7 weeks post-inoculation and 1 mouse at 4 weeks post-inoculation) and *nef*-dimerization defective-Y115D (2 mice at 7 weeks post-inoculation and 1 mouse at 6 weeks post-inoculation) at 4-7 weeks sacrifice-timepoints.

Several studies have demonstrated that HIV Nef promote NFAT-dependent signaling, which inturn promote HIV replication (Abbas and Herbein, 2013; Manninen et al., 2001; Tuosto et al., 2003). IPA of human spleen xenograft-gene expression demonstrates elevated NFAT-dependent signaling in the wild-type HIV group compared to the mock group (Figure 8A). Importantly, IPA of human spleen xenograft-gene expression demonstrates reduced NFAT-dependent signaling in *nef*-deleted HIV group compared to wild-type HIV group (Figure 8B), which suggests that Nef is modulating immune signaling in the human secondary lymphoid tissue (spleen xenograft) in BLTS-humanized mice.

## Discussion

The HIV accessory protein Nef has been implicated in HIV replication, immune dysregulation, and immunodeficiency (Boasso et al., 2009; Chirmule et al., 1994; Foster and Garcia, 2008; Gorry et al., 2007; Poe and Smithgall, 2009; Trible et al., 2006a). Although humans and monkeys infected with *nef*-defective HIV and SIV, respectively, have provided insights into the pathogenic and immune inhibitory role of Nef (Daniel et al., 1992; Dyer et al., 1999; Learmont et al., 1999), humanized rodents provide the ideal small animal model for *in vivo* mechanistic studies of HIV-immune system interactions (Agarwal et al., 2020; Denton and Garcia, 2011; Samal et al., 2018). Previous studies in the BLT-humanized mouse model confirmed the role of HIV Nef in mediating HIV-induced CD4+ T cell depletion (Zou et al., 2012). However, *nef*-defective HIV viremia in the BLT-humanized mouse model was only delayed for a few weeks (at low doses of inoculum) or reduced (at high doses of inoculum) (Zou et al., 2012). The delayed *nef*-defective HIV viremia in the BLT-humanized mouse model suggest that a humanized mouse model with an improved human immune system containing both primary and secondary human lymphoid tissue xenografts could completely mitigate *nef*-defective HIV viremia.

We recently incorporated a human secondary lymphoid tissue (the spleen xenograft) in the BLT-humanized mouse model to create the BLTS-humanized mouse model. The BLTS-humanized mouse model enabled T cell development and education in the human thymic microenvironment (Kondo et al., 2019) and antigen-specific T cell expansion in the human splenic microenvironment (Lewis et al., 2019). Furthermore, the BLTS-humanized mouse model is reconstituted with various human innate and adaptive immune cells, including macrophages, T, and B cells (Samal et al., 2018). Here, we demonstrate the functionality of the antigen-presenting cells and T lymphocytes cells in BLTS-humanized mice, suggesting that those cells can enable an effective antiviral T cell immune response. We previously demonstrated that the BLTS-humanized mouse model supports HIV replication (Samal et al., 2018). HIV viremia kinetics in BLTS-humanized mice mimic patterns in HIV-infected adult humans, such as the partial control of viremia after peak viremia 2-weeks post-infection (Samal et al., 2018). Here, we employed the BLTS-humanized mouse model to determine the role of Nef expression and homodimerization in HIV viremia and associated immune dysregulation.

First, we demonstrated that *nef*-defective HIV and wild-type HIV exhibit similar infectivity and replication at high inocula in a tissue culture model (activated CD4+ T cells) and an *in vivo* model (PBL-humanized mice containing activated CD4+ T cells) of HIV infection and replication. In contrast, *nef*-deleted HIV and wild-type HIV exhibit divergent viremia outcomes at high inocula in BLTS-humanized mice; the *nef*-deleted HIV group remains aviremic for 20-weeks post-inoculation, while the wild-type HIV group exhibits high viremia by 2 weeks post-infection followed by a plateau in viral load up to 6 weeks. Furthermore, HIV viremia was associated with CD4+ T cell depletion in the blood. Both *nef*-defective HIV and wild-type HIV elicited an initial burst of human cytokine responses in the BLTS-humanized mice, although the burst was higher with wild-type HIV infection. This elevated cytokine response in wild-type HIV infection is likely due to the high viremia, compared to the aviremic, *nef*-defective HIV. Notably, the elevation of human cytokine levels suggests a human immune response to both *nef*-defective HIV and wild-type HIV, with an adequate response elicited by *nef*-defective HIV and an ineffective response elicited by wild-type HIV.

Nef enhances of HIV infectivity, replication, and immune dysregulation in tissue culture models (Foster and Garcia, 2008; Poe and Smithgall, 2009). Previous studies have demonstrated that homodimerization is essential for Nef-mediated enhancement of HIV infectivity, replication, and immune dysregulation in tissue culture models (Foster and Garcia, 2008; Poe and Smithgall, 2009). Here, we show that Nef homodimers enhances HIV viremia and associated T cell activation (i.e., elevated CD25 and HLA-DR expression) and expression of a checkpoint inhibitor (i.e., high PD1 expression) and general immune dysregulation in a small animal model with a robust human-immune system. The level of disruption in Nef dimerization activity was consistent with the viremia outcomes. Additionally, gene expression analysis of the human secondary lymphoid tissue (human spleen xenograft) suggests that functional Nef induces NFAT-dependant signaling, a well-established Nef-signaling pathway target in tissue culture models (Abbas and Herbein, 2013; Manninen et al., 2001; Tuosto et al., 2003). Gene expression analysis of the human secondary lymphoid tissue (human spleen xenograft) also suggests that functional Nef promotes the expression of a subset of non-receptor tyrosine kinases (ABL1 and SYK), induces general immune dysregulation (including B cell dysregulation), and impairs the expression of anti-viral factors *in vivo*.

In summary, we report for the first time that Nef expression and dimerization mediate enhancement of HIV viremia and promotes T cell activation, checkpoint inhibitor expression, and general immune dysregulation *in vivo* in the BLTS-humanized mouse model. T cell hyperactivation, exhaustion, and widespread immune dysregulation are believed to be significant drivers in maintaining the HIV reservoir. Furthermore, recent evidence demonstrates that Nef is expressed in HIV-infected individuals receiving highly active-ART (Ferdin et al., 2018; Wang et al., 2015), suggesting Nef could play a significant role in maintaining the HIV reservoir by supporting low-level infection and replication and dysregulating the anti-viral immune response. The BLTS-humanized mouse model supports an ART-mediated HIV reservoir (Samal et al., 2018), thus providing an ideal model for investigating the impact of Nef-inhibitors (including inhibitors targeting Nef dimerization) in HIV cure strategies.

## Supplementary Figures and Legends

**Supplementary Figure 1.**
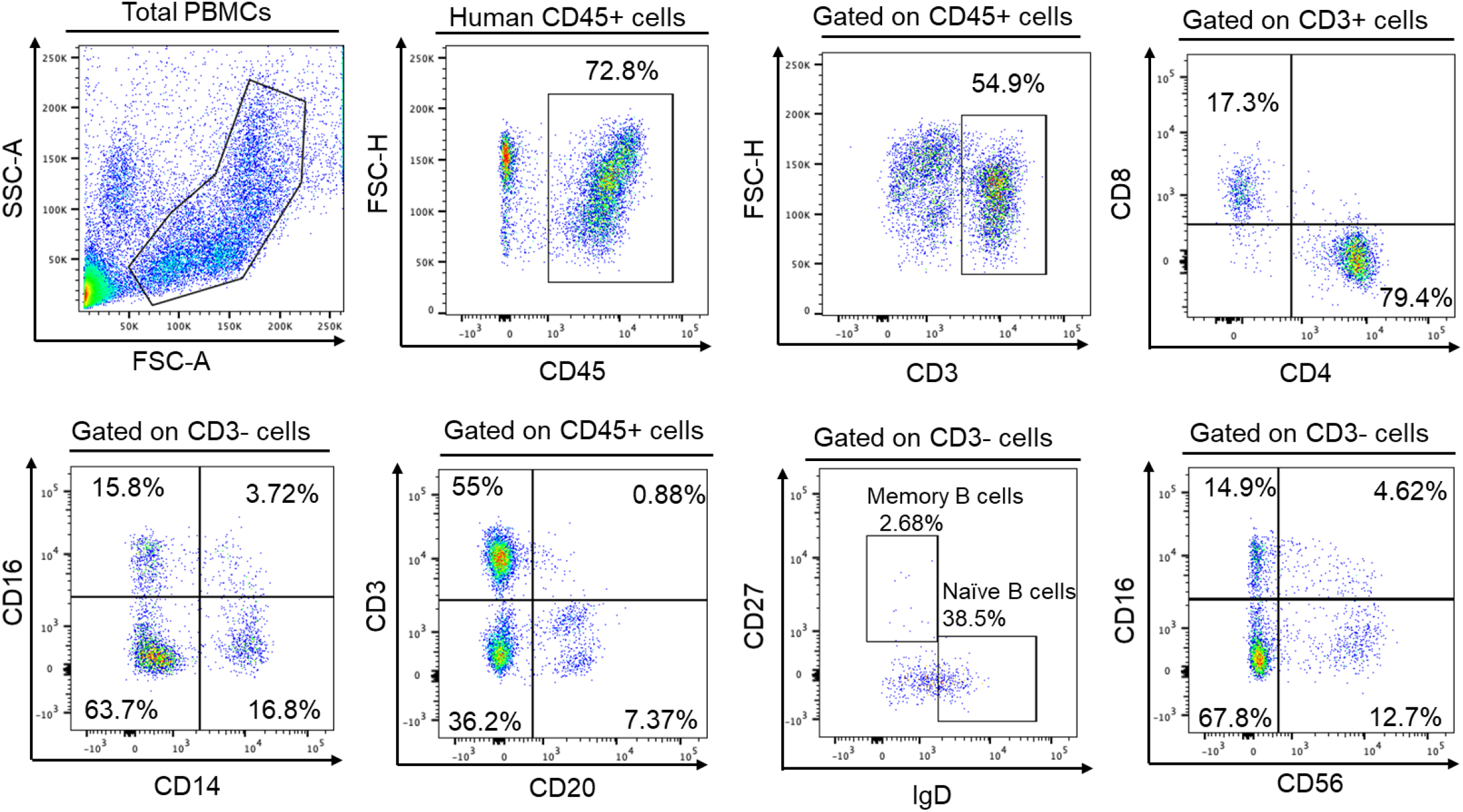
Human peripheral blood cell reconstitution in BLTS-humanized mouse model. Flow cytometry analysis of the blood from representative BLTS-humanized mouse demonstrate the presence of human leukocytes (CD45+ cells), CD3+ T cells (CD4+ and CD8+ cells), monocytes (CD14+ CD16+ cells), (CD20+CD3-) B cells (memory and naive) and NK cells (CD56-CD16+).

**Supplementary Figure 2.**
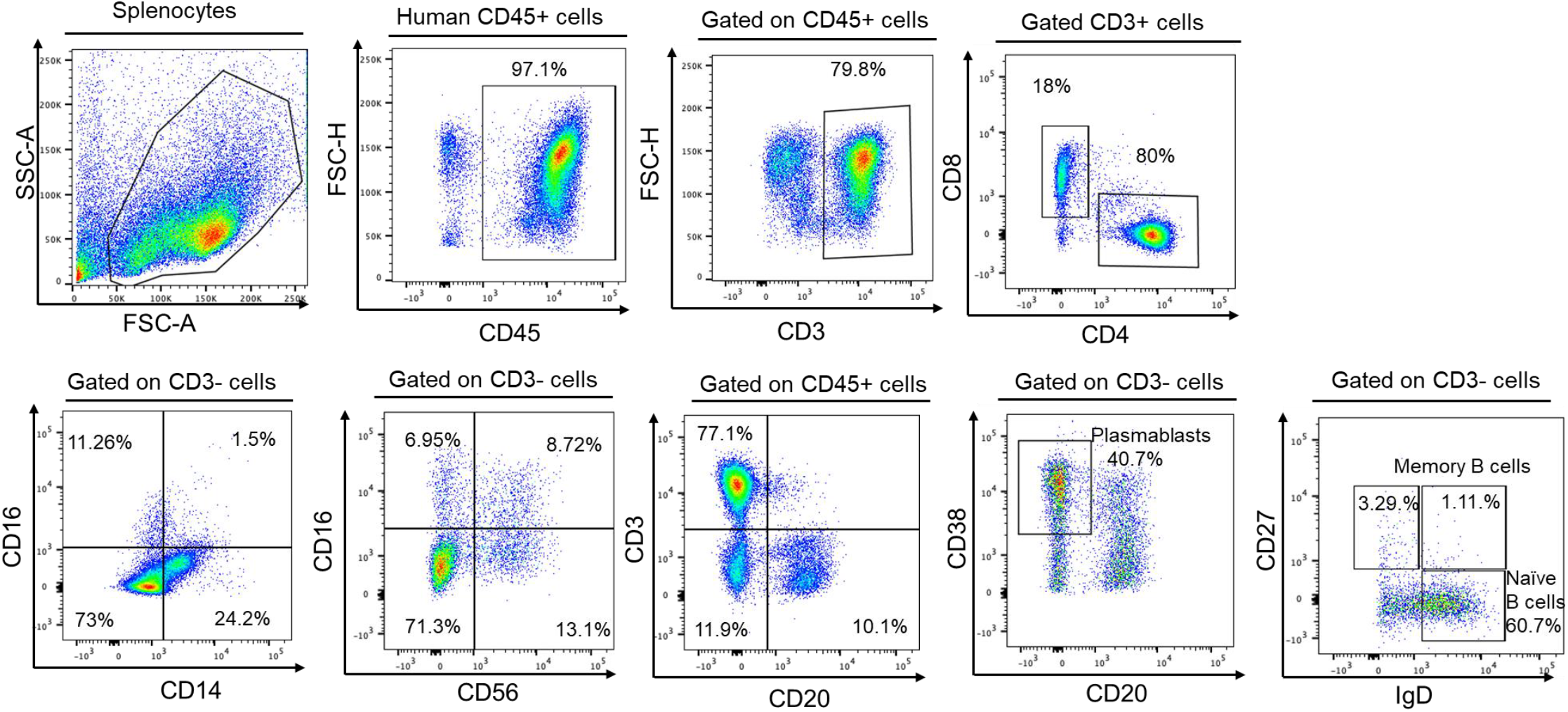
Human immune cell reconstitution in the human spleen in the BLTS-humanized mouse model. Flow cytometry analysis of the human splenocytes from a representative BLTS-humanized mouse demonstrate the presence of human leukocytes (CD45+ cells), T cells (CD4+ and CD8+ cells), monocytes (CD14+ CD16+ cells), NK cells and B cells (including plasmablasts, naive and memory B cells).

**Supplementary Figure 3.**
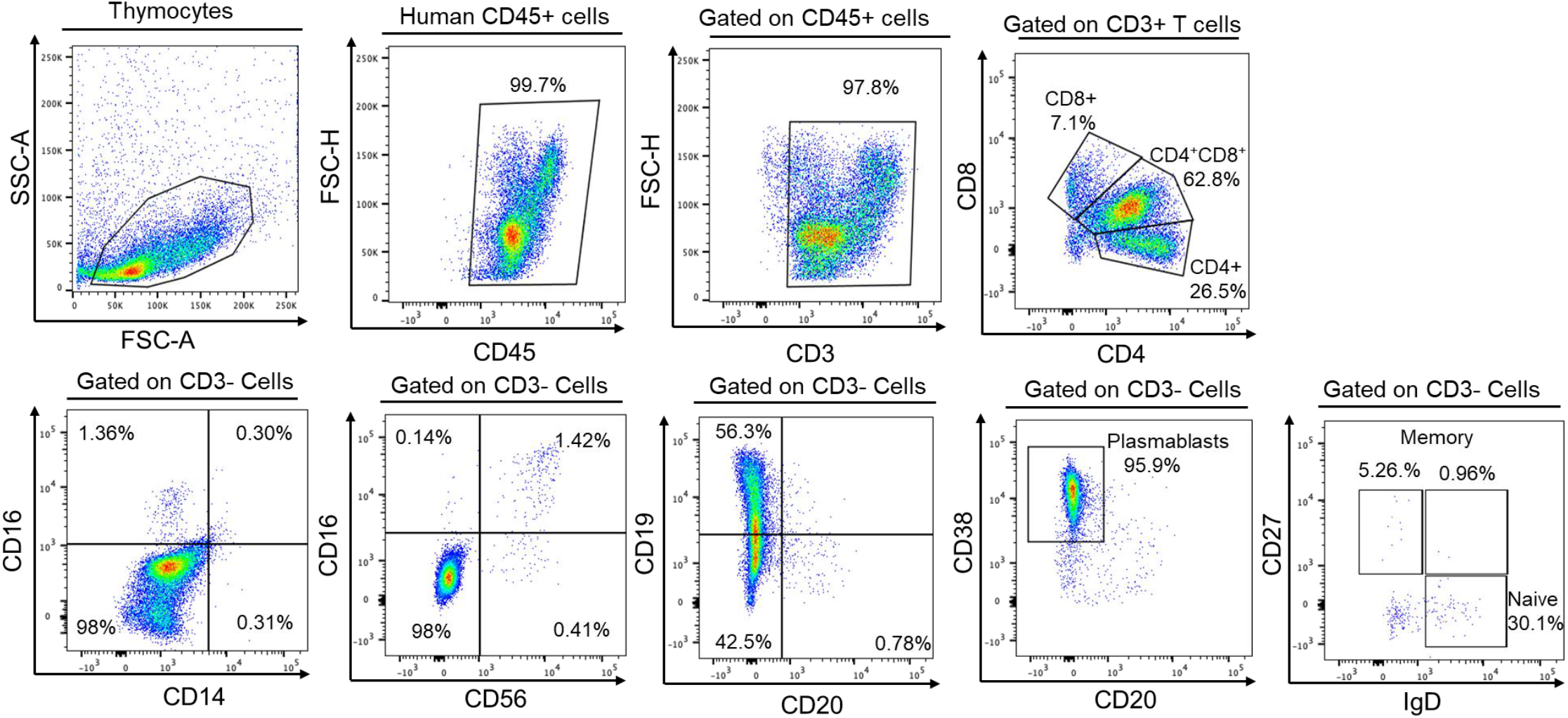
Human immune cell reconstitution in the human thymus in the BLTS-humanized mouse model. Flow cytometry analysis of the human thymocytes from a representative BLTS-humanized mouse demonstrate the presence of human leukocytes (CD45+ cells), including T cells (CD4+ and CD8+ cells) and very low levels of human monocytes (CD14+ CD16+ cells), NK cells and B cells (including, plasmablasts, naive and memory B cells).

**Supplementary Figure 4.**
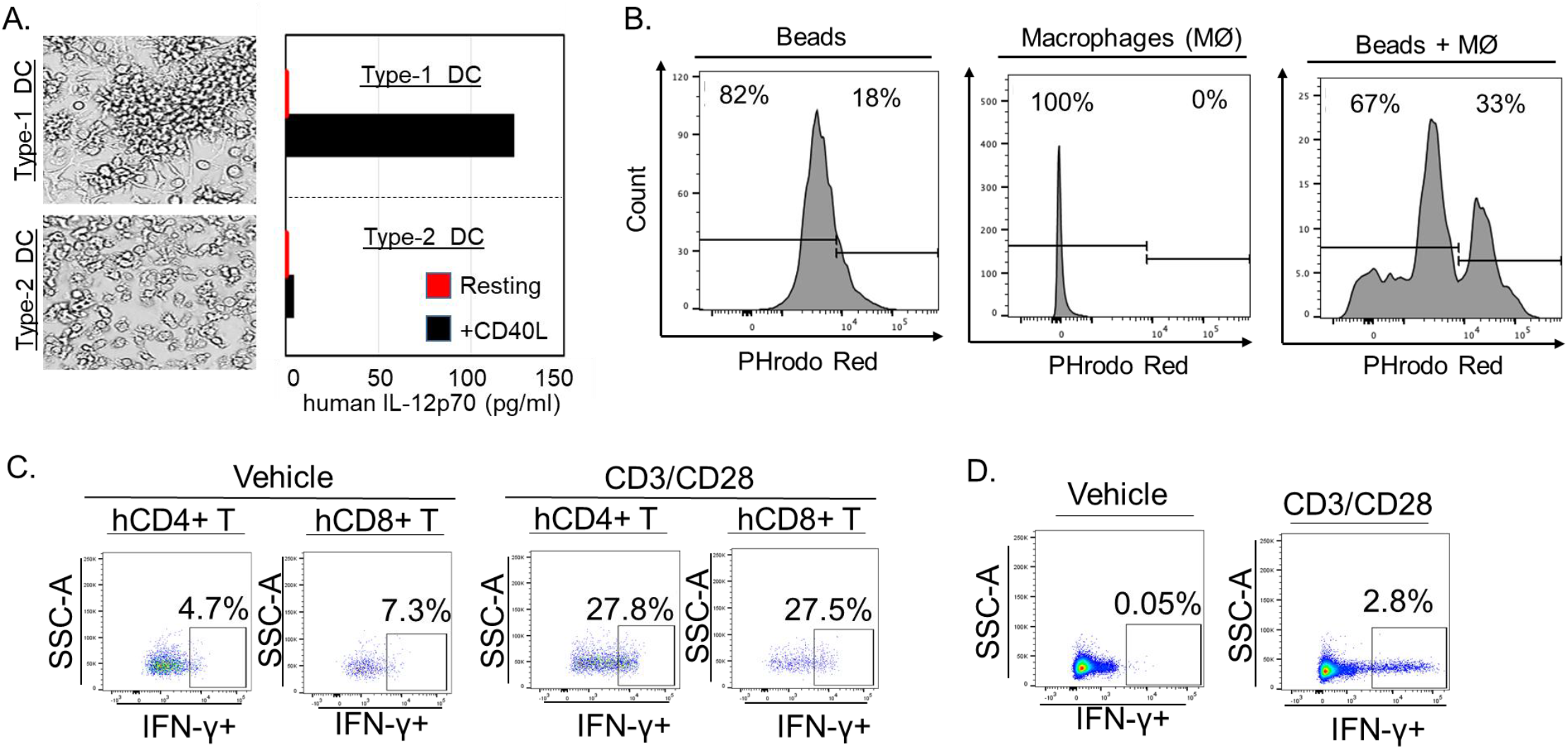
The functionality of human antigen presenting cells and T cells in the BLTS-humanized mouse model. (A) Human CD34+ cells were differentiated into Type-1 and Type-2 dendritic cells (DC), and subsequently stimulated with CD40 ligand or vehicle, and human IL12p70 was measured. (B) Human CD163+ macrophages were immunoselected from human splenocytes and cultured with or without pHrodo™ Red E. coli BioParticles™ (ThernoFisher), and phagocytosis was measured using flow cytometry. (C-D) Human CD3+ T cells were selected from BLTS-humanized mice (human splenocytes) (C) and adult human PBMCs and stimulated with CD3/28 beads, and human IFN-γ secretion was measured via flow cytometry.

**Supplementary Figure 5.**
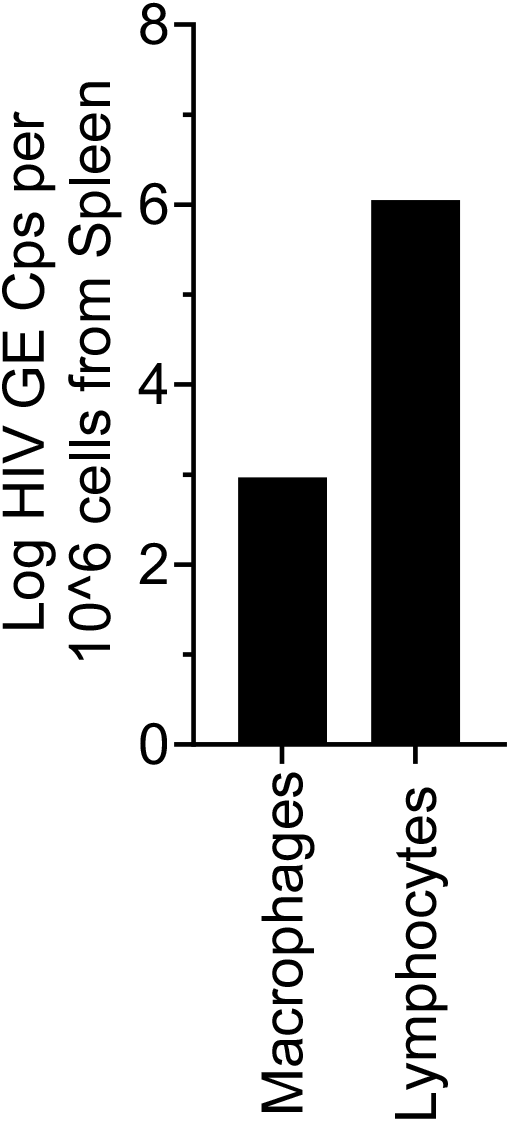
Human CD163+ macrophages and lymphocytes support HIV replication in the human spleen xenograft. Human CD163+ macrophages and CD163-splenocytes were isolated via immunoselection (using human CD163+ immunomagnetic beads) from human splenocytes in BLTS-humanized infected with HIV (NL4) for 2 weeks. Total RNA was isolated from both CD163+ cells (macrophages) and CD163-cells (lymphocytes) and HIV viral load was measured using qPCR.

**Supplementary Figure 6.**
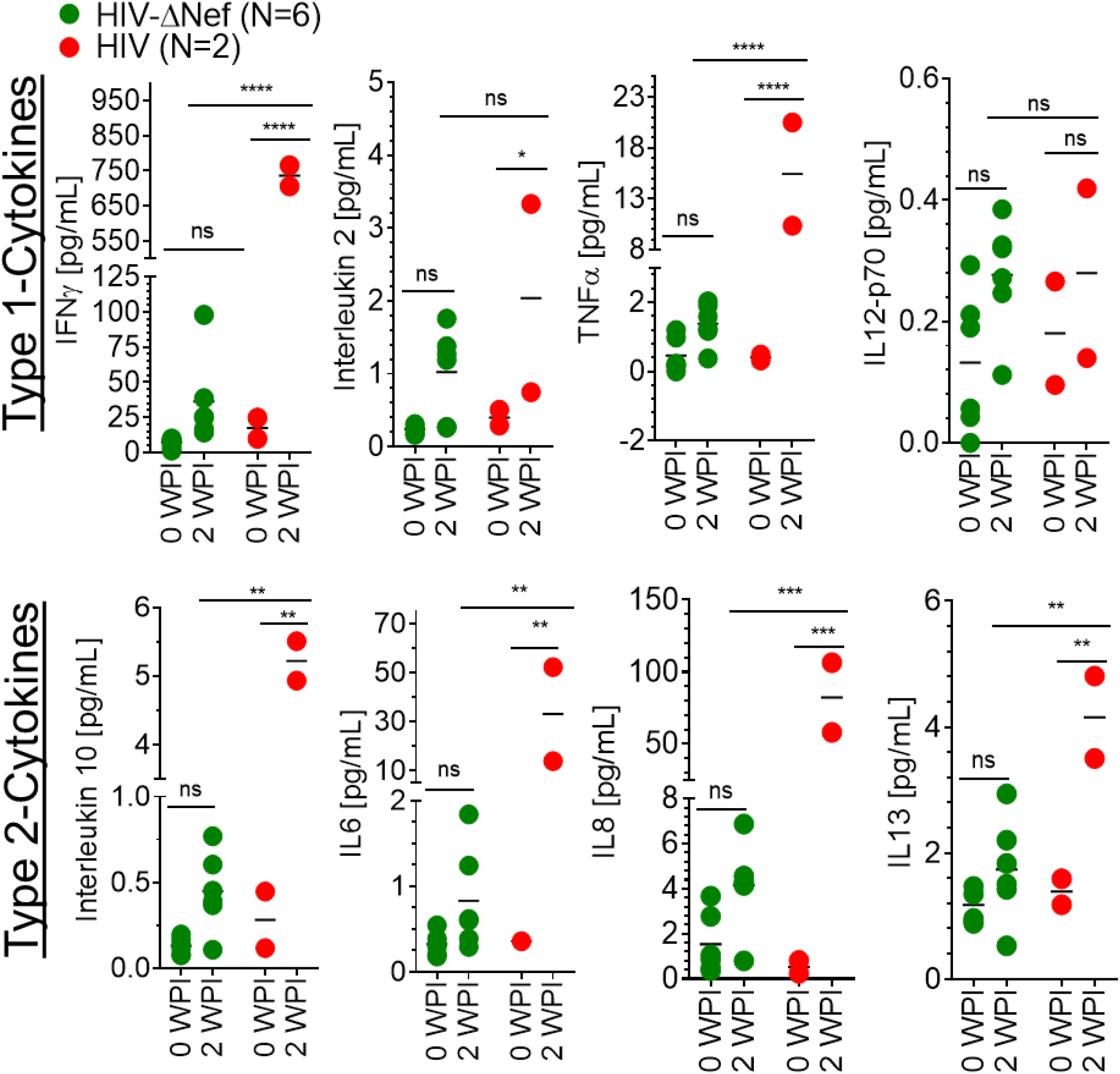
The human-immune system in BLTS-humanized mice mediate a cytokine response to *nef*-deleted HIV and wildtype HIV transmission. Analysis of the human cytokine levels before and after inoculation with wild-type HIV and *nef*-deleted HIV (10 ng HIVp24 ≈ 1×10^6 infectious units per mouse) in BLTS-humanized mice (N=2-6 mice per group) demonstrate that both viruses induces Th1 and Th2 cytokines albeit, relatively higher levels with wild-type HIV replication. N=2-3 mice per group. ns = P>0.05, * = P≤ 0.05, ** = P≤ 0.01, *** = P≤ 0.001.

**Supplementary Figure 7.**
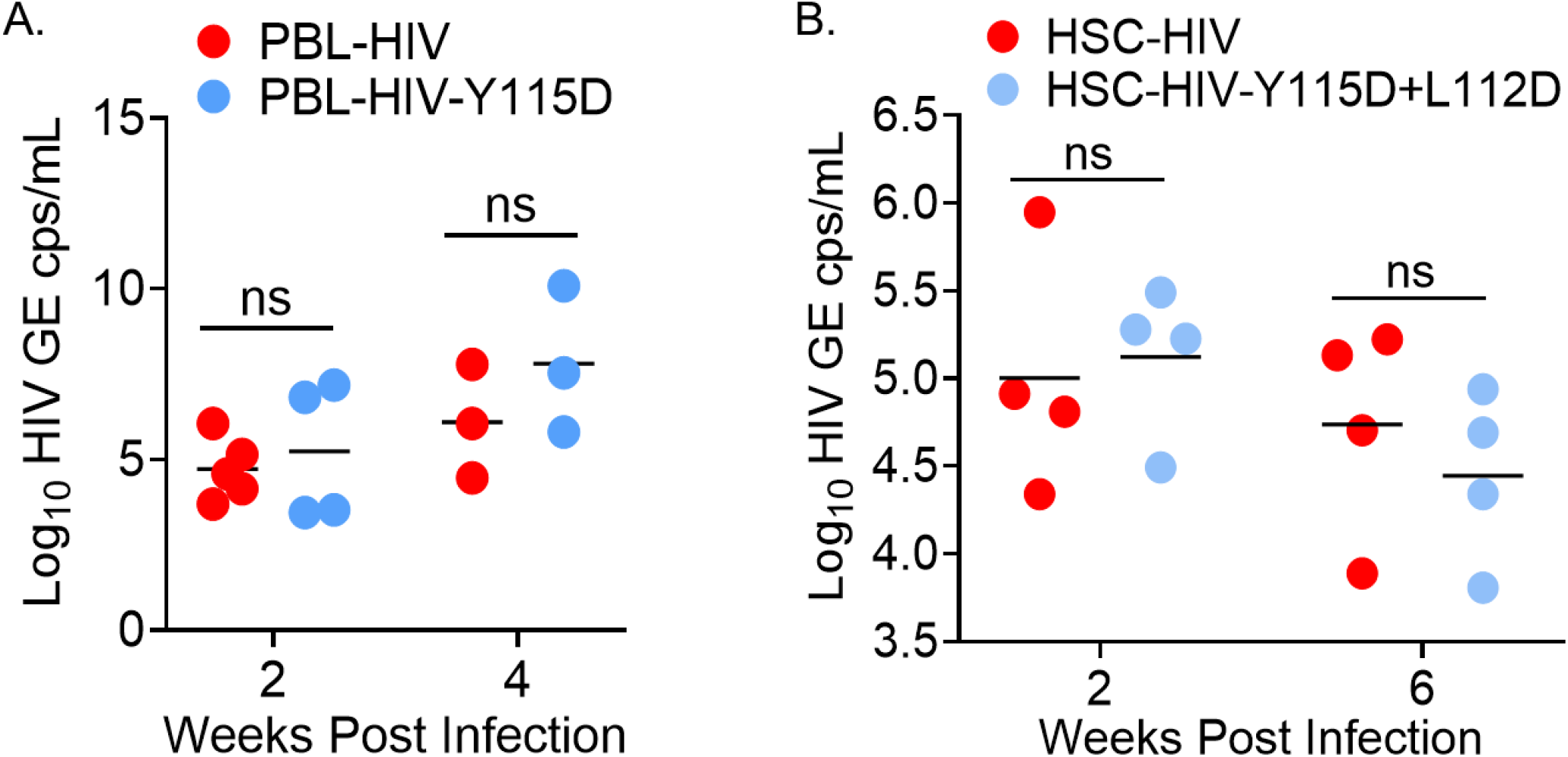
Nef dimerization-defective HIV is replication competent *in vivo*. (A) The infectivity and replication (HIV genome copies/milliliter-GE cps/mL) of wild-type HIV and *nef*-dimerization defective HIV (A. Y115D and B. Y115D+L112D mutant; 2000 Median Tissue Culture Infectious Dose (TCID50) per mouse) in peripheral blood lymphocytes (PBL)-humanized mice and hematopoietic stem cell (HSC)–humanized mice (HSC-Mice) are comparable. N=3-5 mice per group. Not significant (ns) = P>0.05.

**Supplementary Figure 8.**
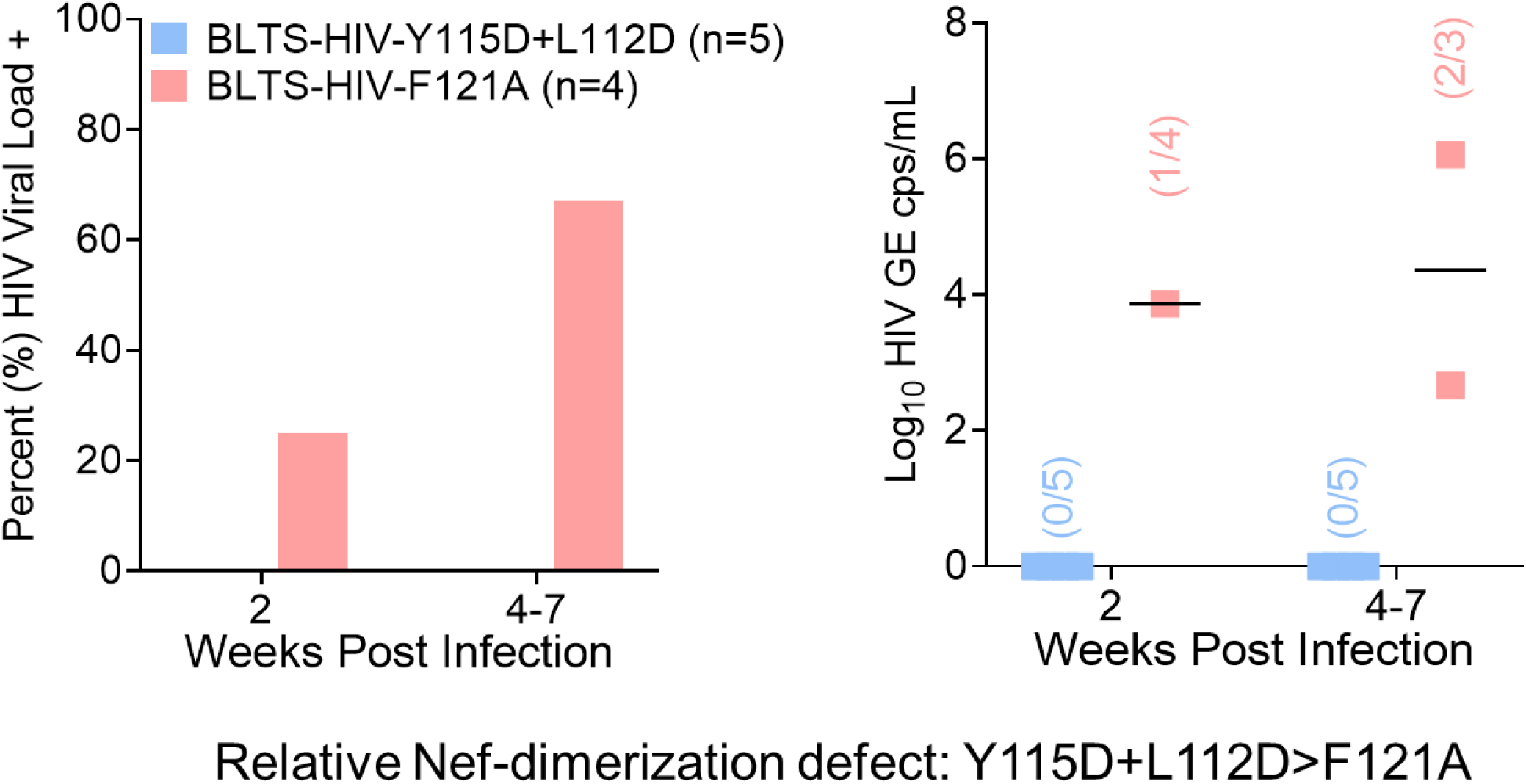
The relative level of Nef-dimerization defect is associated with HIV viremia in the BLTS-humanized mouse model. BLTS-humanized mice (BLTS-Mice) exhibit immune-mediated viremic control of a HIV strain with a higher level of defect in Nef dimerization (Y115D+L112D mutant; 10 ng HIVp24 per mouse) as compared to a HIV strain with a lower level of defect in Nef dimerization (F121A mutant; 10 ng HIVp24 per mouse), which exhibited viremia in most of the mice (67%) at the sacrificed timepoint. N=4-5 mice per group. For BLTS-humanized mice groups, Nef dimerization-defective-Y115D+L112D (5 mice at 6 weeks post-inoculation) and Nef dimerization-defective-F121A (1 mouse at 7 weeks post-inoculation and 2 mice at 5 weeks post-inoculation). Not significant (ns) = P>0.05.

**Supplementary Figure 9.**
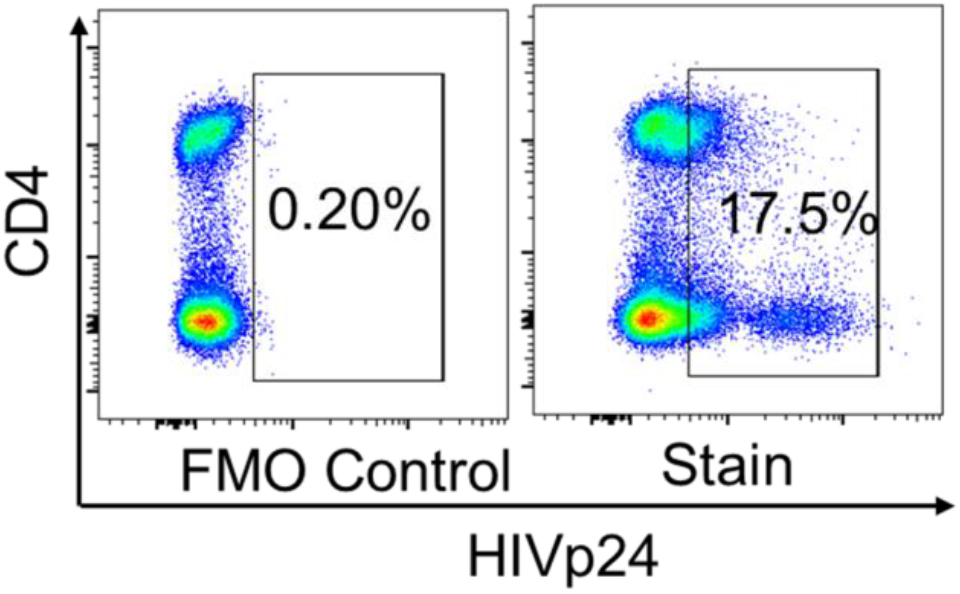
Anti-HIVp24 antibody detection of HIV-infected human PBMCs-derived CD4+ T cells. CD4+ T cells were isolated from healthy human PBMCs using immunomagnetic selection, activated with Anti-CD3/CD28 for three days and subsequently infected with HIV (at multiplicity of infection = one). Ten days post infection, CD4+ T cells were harvested and stained for HIV-p24. HIV-p24 levels in CD4 T cells was analyzed by flow cytometry. HIV-infected human CD4+ T cells were determined based on comparison to fluorescent minus control (FMO).

**Supplementary Figure 10.**
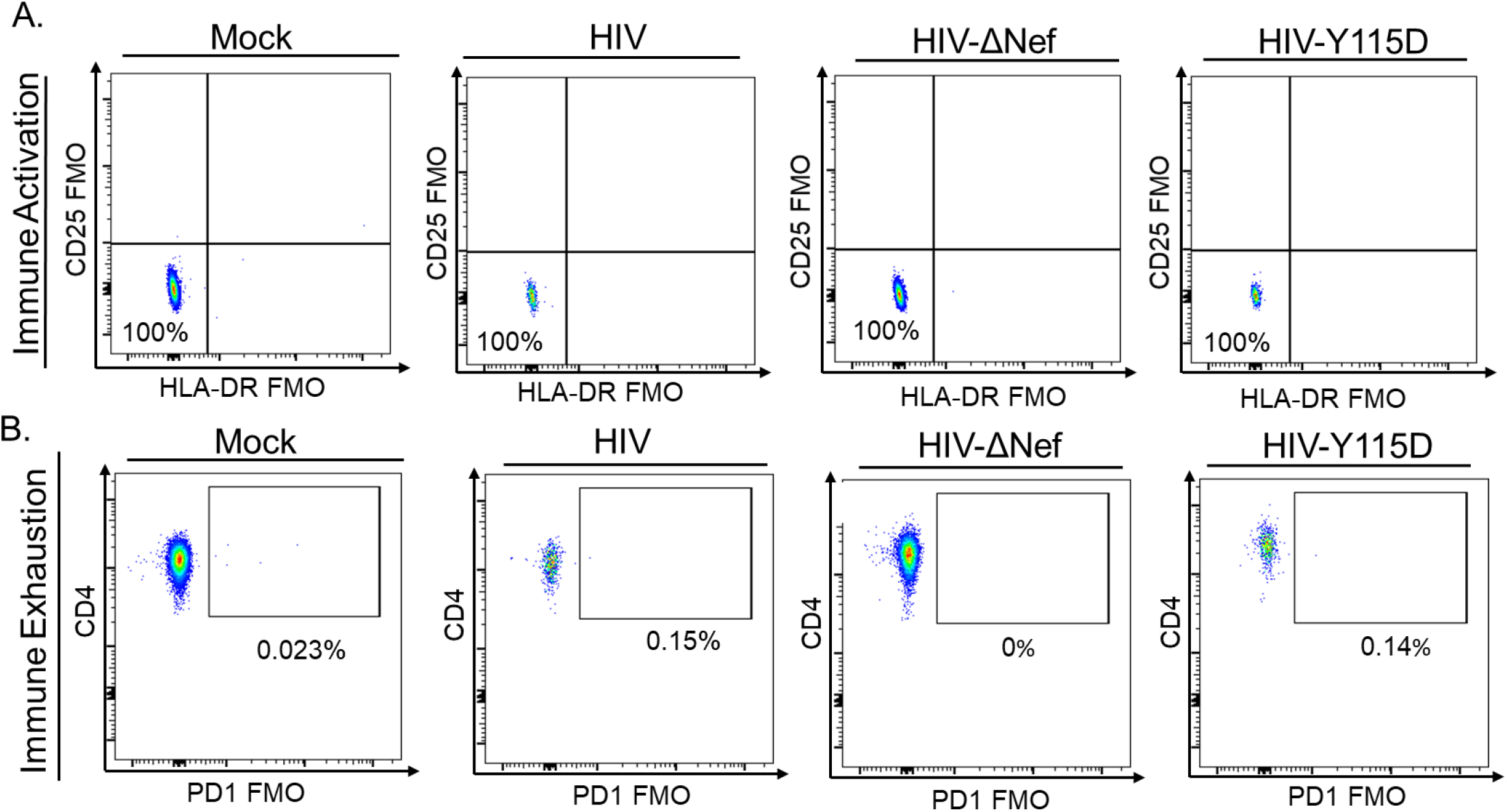
Fluorescent minus staining control for flow cytometry gating of T cell activation and checkpoint inhibitor expression. (A, B) Fluorescent minus staining control-flow cytometry analysis of human CD4+ T cells in the blood in BLTS-humanized mice validates the gating for T cell activation (CD25 FMO plus HLA-DR FMO) (A) and checkpoint inhibitor expression (PD1-FMO) (B).

## Methods

### Virus production

The dimerization mutant constructs used in this study are based on the HIV NL4-3 backbone in which part of the NL4-3 Nef ORF is replaced with that of Nef from the closely related HIV-1 B-clade isolate, SF2. HIV-1 stocks were produced in 293T cells (ATCC; CRL-3216) after transfection and amplified in the T-cell line MT2 (AIDS Reagent Program) (Li et al., 2020; Poe and Smithgall, 2009). Viral titers were quantified by HIV-1 p24 AlphaLISA assay (PerkinElmer; AL291F) according to the manufacturer’s protocol.

### Tissue culture assays

TZM-bl cells and Dynabeads™ Human T-Activator CD3/CD28 (Life Technologies Cat# 11131D) treated human PBMCs-derived CD4+ T cells (via immunomagnetic selection) from an HIV negative donor (10,000 cells per well) were added to separate 96-well plates in RPMI medium 1640-based growth medium (100 ul volume) and cultured overnight. The cells were inoculated with 0.1 ng p24 inoculum (100 ul) of wild-type, nef-deleted, or dimerization defective HIV-1 (NL4-3 strain) and incubated for the indicated time (2, 4, and 6 days). For measuring HIV infectivity and replication in TZM-bl cells, culture supernatant was removed from each well and Beta-Glo® reagent (Promega, Madison, WI) was added and the chemiluminescence activity (Relative Light Units-RLU) was subsequently (after 1 hour) measured using a luminometer following manufacturer’s recommended protocol. For measuring HIV infectivity and replication in human CD4+ T cells, culture supernatant was removed and measured using qPCR (see method below).

### Construction of human immune system/cells-humanized mice

Male and female NOD.Cg-Prkdc^scid^ Il2rg^tm1Wjl^/SzJ (NSG) mice were obtained from the Jackson Laboratory (Stock No: 005557) and bred in the Division of Laboratory Animal Resources (DLAR) facility at the University of Pittsburgh. Human fetal tissues and human peripheral blood were obtained from the Health Sciences Tissue Bank at the University of Pittsburgh or Advance Bioscience Resources and the Central Blood Bank respectively, under approved IRBs. Human tissues and cells were handled and processed under biosafety level 2 conditions. Male and female BLTS, HSC and PBL-humanized mice were generated as previously reported (Samal et al., 2018; Skelton et al., 2018). Briefly, adult NSG mice were sub-lethally irradiated and anesthetized. For BLTS-humanized mice, ∼1-mm^2^ fragments of autologous human fetal thymus, liver and spleen were implanted under the kidney capsule. CD34+ hematopoietic progenitor cells purified from the fetal liver of the same donor were injected intravenously (via retroorbital route) following implantation of the lymphoid tissues. For HSC-humanized mice, CD34+ hematopoietic progenitor cells purified from the fetal liver were injected intravenously (via retroorbital route). For PBL-humanized mice, 20×10^6 PBMCs were transplanted intraperitoneally and immune cells (CD4+ T cells) were allowed to expand over 2 weeks. All mice were housed under specific-pathogen free conditions and fed irradiated chow and autoclaved water. Human immune cell engraftment in BLTS and HSC-humanized mice was detected by flow cytometry at 10-12 weeks after transplantation.

### Generation of monocyte-derived DC

Bone marrow cells were harvested from humanized mice and performed ACK lysis to get rid of RBCs. Then 1 × 10^6^ cells were cultured for 7 days in the presence of 100ng/ml of recombinant human stem cell factor (cat# 255-SC-010),100 ng/mL Flt3-ligand (Cat #-308-FK-005), 50 ng/mL of recombinant human GM-CSF (Sanofi-aventis Cat# NAC2004–5843-01) and 2.5 ng/mL of tumor necrosis factor (TNF)-α (25 ng/mL; R&D Systems Cat# 210-TA). Next, immature DCs were generated from bone marrow cells and cultured for 7 days in Iscove’s Modified Dulbecco’s Media (IMDM; Gibco Cat# 12440-053) containing 10% fetal bovine serum Atlanta biologicals Cat# S12450H) and 0.5% gentamicin (Gibco Cat# 15710-064) in the presence of granulocyte-monocyte colony-stimulating factor (GM-CSF; 1000 IU/mL; Sanofi-aventis Cat# NAC2004–5843-01) and interleukin-4 (IL-4; 1000 IU/mL; R&D Systems Cat# 204-1 L). Mature, high IL-12p70-producing Type 1 DC and IL-12p70 deficient, prostaglandin E2-treated DC (Type 2-DC) were generated as previously described (29) by exposure of immature DC cultures at day 5 for 48 h to a cocktail of maturation factors containing either interferon (IFN)-α (1000 U/mL; Schering Corporation Cat# NDC:0085–1110-01), IFN-γ (1000 U/mL; R&D Systems Cat# 285-1F), IL-1β (10 ng/mL; R&D Systems #201-LB), tumor necrosis factor (TNF)-α (25 ng/mL; R&D Systems Cat# 210-TA), and polyinosinic:polycytidylic acid (20 ng/mL; Sigma-Aldrich Cat# P9582-5MG), or IL-1β (10 ng/mL), TNF-α (25 ng/mL), IL-6 (1000 U/mL; R&D Systems Cat# 206–1 L), and PGE2 (2 μM; Sigma-Aldrich Cat# P6532-1MG), respectively. Differentiation of DC was confirmed by assessing the DC-specific markers such as CD83, CD86, Sig 1, OX40L, CCR-5 by flow cytometry.

### Functional characterization of differentially matured DCs

To test the IL12p70-producing capacity of DC, they were harvested, washed, and plated in flat-bottom 96-well plates at 2 × 104 cells/well. To mimic the interaction with CD40L-expressing Th cells, recombinant human CD40 ligand was added to the culture. Supernatants were collected after 24 h and tested for the presence of IL-12p70 by ELISA.

### Functional characterization of T cells

Immunomagnetically (human CD3+ immunomagnetic beads) selected human splenic T cells from BLTS-humanized mice and human T cells from PBMCs were seeded at 8×10^4 cells per well (96-well plate) and stimulated with Dynabeads™ Human T-Activator CD3/CD28 (Life Technologies Cat# 11131D) and PBS was used as vehicle control. The cells were stimulated with beads at a ratio of 1:1. T cells were harvested after 24 hr and used directly for flow cytometry analysis of the human IFNg levels (T cell activation marker).

### Functional characterization of Macrophages

Humanized mice were euthanized and human splenocytes were isolated from spleen tissue by mechanical grinding. CD163+ splenic macrophages were isolated using Milteny biotech human CD163 MicroBead Kit (cat # 130-124-420). 100,000 macrophages were plated in 96 well plate and incubated in a humidified CO2 incubator for 1 hr. After cells adhered, the culture medium was replaced with 100 μL of the prepared pHrodo™ Red E. coli BioParticles (Cat #-P35361). Next, microplate was transferred to an incubator warmed to 37°C for 1– 2 hours to allow phagocytosis and acidification to reach their maximum. Fluorescence emitted by the pHrodo BioParticle phagocytosed macrophages was analyzed by the flow cytometry.

### Flow cytometry analysis of immune cells

Peripheral blood cells and lymphoid tissue cells were analyzed using flow cytometry. Briefly, peripheral blood was collected from humanized mice and mixed with 20 mM Ethylenediaminetetraacetic acid (EDTA) at a 1:1 ratio. Single cell preparation of blood cells or lymphoid tissue cells were prepared via red blood cells lysis using Ammonium-Chloride-Potassium (ACK) buffer. Single-cell suspensions were stained with a LIVE/DEAD Fixable Aqua Dead Cell Stain Kit (ThermoFisher Scientific), fluorochrome-conjugated antibodies (anti-mouse CD45-BioLegend Cat. No. 103126, anti-human CD45-BioLegend Cat. No. 304014, anti-human CD3-BioLegend Cat. No. 300312, anti-human CD4-BioLegend Cat. No. 317410, anti-human CD8-BioLegend Cat. No. 300906, anti-human CD19-BioLegend Cat. No. 302232, anti-human PD1, BD, cat. No. 562516, anti-human HLA-DR, BD, cat. No. 562304, anti-human CD25, BD, cat. No. 740397, Anti-human CD56 clone N901; Beckman Coulter, CD16-PerCP-Cy5.5 Cat. No. 302027 BD Pharmingen,anti-human CD69, BD, cat. No. 562645, Anti-human CD14 Cat. No. 301819, IFN-γ, BD Biosciences Cat. No. 557995, Anti-HIV-1 p24, KC57-FITC, Beckman Coulter Cat. No. 6604665), fixed, and analyzed on a BD LSRFortessa™ cell analyzer - flow cytometer (BD Biosciences). Data were analyzed using FlowJo software (Dako). Briefly, leukocytes were selected based on forward and side scatter. Single cell and live leukocytes were selected for further analysis of the percentage of human leukocytes (anti-human CD45+, hCD45+) and mouse leukocytes (anti-mouse CD45+, mCD45+) in the peripheral blood. Subsequent analysis of the various human lymphocyte populations and subsets were gated on human leukocytes.

### HIV infection of human immune system-humanized mice

Humanized mice with stable human leukocyte reconstitution were anesthetized and inoculated with HIV-1 retro-orbitally.

### HIV-1 genomic RNA detection

Total RNA was purified from plasma using RNA-Bee (AMSBIO). The RNA was then reverse-transcribed using TaqMan® Reverse Transcription Reagents (Life Technologies) and quantitatively detected by real-time PCR using the TaqMan® Universal PCR Master Mix (Life Technologies) with primers (forward primer - 5’ CCCATGTTTTCAGCATTATCAGAA 3’ and reverse primer-5’ CCACTGTGTTTAGCATGGTGTTTAA 3’) and detection probe targeting HIV Gag gene (5’-AGCCACCCCACAAGA-3’) (Biswas et al., 2012). The assay sensitivity/cutoff is 10 copies/mL (Biswas et al., 2012).

### Cytokine assay

Human cytokine levels were detected in pre and post-infection samples using the V-PLEX Assay and following the manufacturer’s recommended procedures (Meso Scale Diagnostics).

### Gene Expression Profiling Using nCounter Analysis

HIV infected, and Mock-inoculated humanized mice were sacrificed at the end of the study and total RNA was isolated from engrafted spleen tissue of humanized mice using Qiagen RNA isolation kit Cat# 74104. Nanostring profiling of host response was performed at the University of Pittsburgh Genomic Core facility using the nCounter Human Immunology v2 Panel, Cat #-XT-CSO-HIM2-12. Total RNA (50ng) was hybridized to reporter and capture probe sets at 65°C for 24 h. Hybridized samples were loaded on the nCounter cartridge and post-hybridization steps and scanning was performed on the nCounter Profiler. RCC files were analyzed using nSolver analysis software (Version 4.0) as per the manufacturer’s protocols. Negative and positive controls included in probe sets were used for background thresholding, and normalizing samples for differences in hybridization or sample input respectively.

### Statistics

Significance levels of data were determined by using Prism8 (GraphPad Software). Data were analyzed using analysis of variance (ANOVA) and Fisher’s exact tests. A P value less than 0.05 was considered significant. The number of animals is specified in each figure legend.

### Study approval

Human fetal liver and thymus (gestational age of 18 to 20 weeks) were obtained from medically or elective indicated termination of pregnancy through Magee-Women’s Hospital of UPMC via the University of Pittsburgh, Health Sciences Tissue Bank. Written informed consent of the maternal donors was obtained in all cases, under the Institutional Review Board (IRB) of the University of Pittsburgh guidelines and federal/state regulations. The use of human fetal organs/cells to construct humanized mice was reviewed by the University’s IRB Office, which has determined that this submission does not constitute human subjects research as defined under federal regulations [45 CFR 46.102(d or f) and 21 CFR 56.102(c), (e), and (l)]. The use of human hematopoietic stem cells was reviewed and approved by the Human Stem Cell Research Oversight (hSCRO) at the University of Pittsburgh. The use of biological agents (i.e. HIV), recombinant DNA and transgenic animals, etc. was reviewed and approved by the Institutional Biosafety Committee (IBC) at the University of Pittsburgh. All animal studies were approved by the Institutional Animal Care and Use Committee at the University of Pittsburgh and were conducted following NIH guidelines for housing and care of laboratory animals.

## Authors contributions

MB, TS and RM conceived and designed experiments in the study. SB, YA, AD, JS, SS, CB, SH, IC and MB performed experiments. MB, SB, RM, SS, and TS analyzed and interpreted the data. MB, SB and YA prepared the manuscript.

## Acknowledgments

This project used the UPMC Hillman Cancer Center and Tissue and Research Pathology/Health Sciences Tissue Bank shared resource, which is supported in part by award P30CA047904. Ingenuity pathway analysis licensed through the Molecular Biology Information Service of the Health Sciences Library System, University of Pittsburgh was used for data analysis. Holly Anne Bilben perform some viral load analysis and Priyanka Talukdar assisted with the Ingenuity pathway analysis. This work was supported by the National Institutes of Health grants AI126995 (to M.B.) and AI057083 and AI152677 (to T.E.S).

